# Phosphatidylinositol Phospholipids Modulate Hepatitis C Virus Core Protein Assembly on Lipid Membranes

**DOI:** 10.1101/2025.05.13.653720

**Authors:** Titas Mandal, Bastian Albrecht, Shorouk Abdelwahed, Jakob Ruickoldt, Peggy Jones, Petra Wendler, Salvatore Chiantia

## Abstract

Hepatitis C virus (HCV) genome codes for various proteins essential to its replication cycle. Among these, the core protein (HCC) forms the capsid via organised multimerization, thereby guiding viral assembly. This process depends on the dynamic localisation of HCC between the endoplasmic reticulum bilayer membrane and the lipid droplet monolayer. While past studies have examined the role of some HCC structural properties and other viral and host proteins in the assembly process, the contribution of lipid molecules was not thoroughly investigated yet. Since specific lipids are upregulated during HCV infection, it is reasonable that these molecules might play an important role for virus fitness and infection progression. In this study, we have addressed this topic and, specifically, how HCC-lipid interactions might impact HCC-HCC interactions. Using *in vitro* models and quantitative fluorescence microscopy techniques, we investigated HCC binding affinity to lipid membranes, multimerization and lateral diffusivity. Our results reveal the effect of HCC interactions with anionic phospholipids (PLs) on the binding to monolayers and bilayers. Furthermore, our analysis shows that PI(4)P enhances HCC multimerization and, therefore, might drive lateral assembly for capsid formation. These results not only provide insights into the lipid-driven regulation of HCV assembly and identify anionic PLs as potential modulators of the viral replications cycle, but also highlight a previously unrecognised role of PL composition in HCC-membrane interaction.

**Significance:** Hepatitis C virus (HCV) remains a major global health challenge and is associated with substantial mortality. In certain regions, HCV infection reaches epidemic proportions. Despite its prevalence, targeted and cost-effective therapeutic options remain limited, largely due to an incomplete understanding of the viral life cycle. A critical - yet poorly characterized - step in this cycle is the assembly of the HCV core (HCC) protein into the viral capsid, which is essential for replication. Here, we use quantitative fluorescence microscopy to investigate the role of specific lipid species in HCC assembly on model membranes. Our findings show that distinct lipid environments modulate HCC assembly and that HCC, in turn, induces measurable changes in the biophysical properties of the host membrane.

## Introduction

Infection by Hepatitis C virus (HCV) is associated to severe liver disease and poses a serious global health challenge, due to the absence of vaccines and broadly accessible targeted medications (1–3). HCV is an enveloped virus with a 9.6 kb positive-sense RNA that codes for various viral proteins, including the Core protein (HCC) (4). HCC forms the structural subunits of the viral capsid, i.e. an icosahedral protective shell between the viral genome and the lipid envelope of the virus (5). Additionally, HCC plays fundamental roles in viral assembly (4–7), as well as hijacking of several host cellular pathways for sustaining the viral replication cycle (8–13).

HCV RNA translates into a polyprotein which produces ten individual viral proteins upon cleavage at specific sites by the host and viral peptidases (11). HCC is derived from the N-terminus of this polyprotein via the action of a host peptidase (11, 14, 15). The 191 amino acids (aa) long nascent HCC is targeted to the endoplasmic reticulum (ER) membrane owing to the presence of the C-terminal signal peptide (SP) domain (20 aa), which also acts as a bilayer transmembrane anchor (4, 16). Besides the SP, HCC includes two more domains: D1 and D2 (17). D1 is a larger (117 aa) hydrophilic domain enriched in basic amino acids. D2 is a shorter (54 aa) and more hydrophobic domain that consists of two amphipathic helices (H1 and H2) connected by a hydrophilic loop (HL) (16–18). Owing to the presence of the basic residues, D1 exhibits diverse interactions with the viral RNA (19–21) and, subsequently, promotes the formation of nucleocapsid (4, 22). Additionally, D1 promotes homotypic ionic interactions that form homo-multimers, necessary for capsid formation (6). The D2 domain, on the other hand, interacts with lipid membranes through its amphipathic character (16, 23). D2 additionally promotes HCC homo-dimer formation through a disulphide bond between the Cysteine residues (Cys128) of each protein (18). Such dimers play a notable role in the viral assembly by establishing the foundation for a stable nucleocapsid (5, 18, 24). As a part of the HCV replication cycle, HCC exhibits an organized dynamic partition between the ER bilayer membrane and lipid droplet (LD) monolayers (14–16, 25, 26). At the ER membrane, SP-peptidase-mediated cleavage of SP gives rise to the mature form of the HCC (171 aa) (16), which is subsequently escorted to the LD surface by host proteins (25, 27). The HCC-LD relationship is of paramount importance for the viral assembly process (15). For instance, HCC starts assembling into highly ordered multimers while on the surface of LDs, marking the first step of the viral assembly (15). Interestingly, arresting this step disrupts the viral life cycle (15) by inhibiting the transfer of HCC from the LD back to the ER (28–30). HCC multimer re-targeting to the ER membrane is actively mediated by other HCV proteins (4, 14, 26), and this translocation has been shown to assist in the late steps of the viral life cycle (31). These include synchronised packaging of the viral RNA with the HCC multimers and other viral proteins to form new viral particles (14, 26), which subsequently transit into the host secretory pathway (4, 22, 30, 32). Interestingly, HCC itself can form virions, independent of other viral proteins (15), indicating the fundamental role of this protein in virion production.

While (protein) host factors involved in HCC trafficking between ER and LDs were studied extensively (14–16, 25, 26), the influence of lipids in determining HCC localization and viral assembly are not well understood. Recent findings highlighted the role of neutral lipids such as triglycerides (TGs) in promoting D2 binding to artificial LDs (ALDs) or membrane bilayer models (i.e., giant unilamellar vesicles (GUVs)) (33). On the other hand, zwitterionic-phospholipids (PLs) such as phosphatidylcholine/ phosphatidylethanolamine were shown to hinder D2 interactions with both monolayers and bilayers (33). Otherwise, no detailed information is available regarding how specific PLs might affect HCC-lipid and HCC-HCC interactions. In this context, it is worth mentioning that, as a part of its host cellular pathway hijacking process, HCV upregulates the synthesis of certain anionic PLs such as phosphatidylinositol phosphates (PIPs) (which are in fact identified as “clinical hallmark” of HCV infection) (34–36). Despite their significant involvement in various stages of the HCV replication cycle (37), the possible contribution of anionic PLs in HCC trafficking and subsequent assembly is not known.

In this study, we clarify the role of anionic PLs in HCV assembly, using *in vitro* approaches. To this aim, we investigated the interaction of fluorescently labelled recombinant HCC (18) with model membranes, such as GUVs and ALDs representing PL bilayers (ER membrane) and LD monolayers, respectively (33, 38, 39). Specifically, we quantified HCC interactions with ALDs and GUVs in terms of binding affinity, multimerization, lateral dynamics and assembly using fluorescence fluctuation microscopy methods, namely line-scan fluorescence correlation spectroscopy (lsFCS) (40, 41). This method was successfully used in the past to study viral protein assembly, *in vivo* localisation and dynamics (41–43). By performing such a systematic analysis of the interactions between HCC and model membrane platforms composed of different PLs, we show that specific anionic PLs strongly influence HCC multimerization at the surface of LD, and, therefore, might play a fundamental role in the assembly of HCV.

## Material and Methods

### 1. Materials

For the preparation of ALDs and GUVs, the detergent n-dodecyl β-D-maltoside (DDM) and the following lipids were purchased from Avanti Polar Lipids (Alabaster, AL, USA): 1,2-dioleoyl-sn-glycero-3-phosphocholine (DOPC), sphingomyelin (Milk, Bovine), 1,2-dioleoyl-sn-glycero-3-phospho-L-serine (DOPS), L-α-phosphatidylinositol-4-phosphate (Brain PI(4)P), L-α-phosphatidylinositol-4,5-bisphosphate (Brain PI(4,5)P2), 1-oleoyl-2-(6-((4,4-difluoro-1,3-dimethyl-5-(4-methoxyphenyl)-4-bora-3a,4a-diaza-s-indacene-2-propionyl)amino)hexanoyl)-sn-glycero-3-phosphocholine (TF-PC) and 1,2-distearoyl-sn-glycero-3-phosphoethanolamine-N-lissamine rhodamine B sulfonyl-ammonium salt (Rh-PE). Glycerine trioleate (Triolein) for the preparation of LDs, Alexa Fluor^®^ 488 dye (AF488) and Alexa Fluor^®^ 555 dye (AF555) for the calibration of excitation beam waist, and streptavidin for GUV chamber coating were purchased from Thermo Fisher Scientific (Waltham, MA, USA). Sucrose for electroformation was purchased from PanReac Applichem GmbH (Darmstadt, Germany). Tween^®^20, Tris-HCL, Tris base, magnesium chloride (MgCl_2_), sodium chloride (NaCl), potassium chloride (KCl), urea, β-mercaptoethanol, Bovine serum albumin (BSA) for GUV chamber coating, disodium phosphate and monosodium phosphate for sodium phosphate buffer (NaP) preparation, and trifluoroacetic acid were purchased from Carl Roth GmbH (Karlsruhe, Germany). Skim milk for blocking buffer, PMSF, and DNase I were purchased from Sigma-Aldrich (St. Louis, Missouri, USA). EDTA-free protease inhibitor cocktail tablet was purchased from Roche Holding AG (Basel, Switzerland).

### 2. Vector expression and protein purification

The plasmid CMV-FLAG-Core-R (catalog no. 22480) for expressing HCC (genotype 1b), was purchased from Addgene (Watertown, MA, USA). A forward primer with Nde1 (TAACATATG**ATGAGCACGAATCCTAAACCTC**) and reverse primer with BamH1 (TTAGGATCC**CAGATTCCCTGTTGCATAGTTC**) cleavage site was designed to amplify a 507-bp nucleotide fragment corresponding to the 169 aa long HCC (18) from the purchased plasmid, using PCR. This amplified segment was then ligated into pET-15b vector (Novagene, catalog no. 70755, Burlington, MA, USA) and transformed into *E.coli* (BL21 AI strain) (Thermo Fisher Scientific, Waltham, MA, USA) for protein expression. N-terminal 6X-Histidine tagged HCC was expressed and purified following the available protocol by Boulant *et al*. (18) with slight modifications.

*E.coli* BL21 AI transformants were incubated in LB medium mixed with 100 µg/mL ampicillin at 37 °C (120 rpm) until the growth culture optical density (at 600 nm) reached ~0.7. Protein expression was induced using L-arabinose and IPTG at final concentrations of 0.2 % (w/v) and 1 mM, respectively, and allowed to grow for a further three hours. Then, the bacterial cells were harvested by centrifugation at 4,248 x g. Following this, the obtained pellet was resuspended in ice-cold Buffer A (25 mM Tris-HCL pH 7.5, 5 mM MgCl_2_, 1 mM PMSF, EDTA-free protease inhibitor cocktail tablet (1 tablet for 0.5 mL), and 100 µg/mL DNase I). Next, the cells in the suspension were lysed using a French pressure cell press (Polytec, Germany) at 1200 psi and centrifuged at 38,759 x g for one hour at 4° C. As expected (18), the expressed HCC was enriched in inclusion bodies and, therefore, the pellet was further processed. The pellet was resuspended in Buffer B (20 mM Tris-HCL pH 8, 500 mM NaCl, 6 M urea, 10 mM β-mercaptoethanol and 0.1 % (w/v) DDM) and sonicated for homogenization. The homogenate was subjected to centrifugation. All centrifugation steps were performed using Sorvall Lynx 4000 centrifuge (Thermo scientific, Waltham, MA, USA). Cell harvesting was performed using a F12-6x500 LEX rotor (Thermo scientific, Waltham, MA, USA) and the remaining centrifugation steps for protein separation were performed using a A27-8x50 rotor (Thermo scientific, Waltham, MA, USA).

The filtered supernatant was loaded over a Ni-NTA-agarose column (Qiagen, Germany) and purified using an FPLC system (Biorad NGC chromatography system, Hercules, California, USA) following the manufacturer’s protocol. All purified fractions were screened by SDS-PAGE (15 % resolving gel) and Coomassie Blue staining. The fractions consisting of HCC (~18 kDa) were isolated and further purified by reverse-phase HPLC (Shimadzu, Japan) using a VYDAC 10 µm C8 column (250 x 4.6 mm) equipped with a 208TP Aquapore guard cartridge (Avantor, Pennsylvania, USA). A linear gradient of acetonitrile in 10% trifluoroacetic acid was used as the mobile phase. All fractions were monitored at 280 nm with an integrated UV-visible detector. The main absorption peaks were screened by SDS-PAGE and silver staining. DDM was added (0.1 % (w/v) for 1 mg/mL protein) to the HCC containing fractions for protein stabilization. Finally, the HCC containing fractions were lyophilized and stored at −20 °C until any further use. Unless mentioned, the lyophilized protein fractions were resuspended in ice cold 20 mM NaP buffer (pH 7.4) when needed. For fluorescence microscopy experiments, protein labelling was performed using the primary amine reactive dye Alexa Fluor^®^ 488 (AF488) following the manufacturer’s protocol. In this regard, a degree of labelling (*DOL*) ~0.2 was reached, as confirmed via optical spectroscopy measurements (data not shown).

Only a minor modification to the protocol was needed for the HCC mutant Δ68 (44), which was purified from the soluble fraction of the bacterial lysate using urea free buffer and labelled following the same protocol.

### 3. PIP strip™ blotting

PIP strips™ (Thermo Fisher Scientific, catalog no. P23751, Waltham, MA, USA) were used for screening the PL binding preference of the purified HCC according to a published protocol (45). For blocking unspecific interactions, each membrane was incubated in Buffer C (137 mM NaCl, 2.7 mM KCl, 19mM Tris base, 0.1 % (v/v) Tween^®^20) + 1 % (w/v) skim milk (blocking buffer) for one hour at room temperature (RT). Subsequently, the blocked strip was incubated overnight with 1 µg/mL unlabelled HCC in 20mM NaP buffer at 4 °C. The incubated strip was then washed three times with Buffer C and subjected to one hour incubation with mouse anti-HCV Core antibody (Thermo Fisher Scientific, catalog no. MA1-080, Waltham, MA, USA) in blocking buffer (1/1000 dilution). The washing step was then repeated, followed by the addition of goat anti-mouse alkaline phosphatase (AP) conjugated secondary antibody (Invitrogen, catalog no. 31320, Waltham, MA, USA) (1/5000 dilution). After one hour incubation with the secondary antibody, the washing step was repeated, and the membrane was treated with the AP substrate kit (BioRad, catalog no. 1706432, Hercules, California, USA) for chromogenic detection of possible protein binding to the PL spots using the manufacturer’s protocol. Images of the developed membrane were acquired using an image scanner (ImageScanner III, GE, Boston, MA, USA). Quantification of protein binding intensities was performed using Fiji software (46).

### 4. Preparation of ALDs

ALDs were employed as *in vitro* lipid monolayer platforms for the quantitative investigation of HCC-lipid interaction. ALDs were prepared using triolein as the oil phase according to an available protocol (47) with slight modifications (39). Firstly, a PL solution mixed with 0.001 molar% (w/v) Rh-PE (or 0.005 molar% TF-PC) in methanol was dried in a microcentrifuge tube under a nitrogen stream. The dried film was dispersed in pre-warmed triolein (1/1000 mass ratio) and the mixture was extensively vortexed for 15 minutes and bath sonicated for 30 minutes. Throughout the manuscript, triolein is referred to as “oil”. The solution was then added to 20 mM NaP buffer of pH 7.4 (1:10 v/v) and sonicated for ten seconds after short mixing, thus obtaining a milky white oil-in-buffer emulsion which contained the ALDs. 75 µL of this ALD emulsion was transferred to an observation chamber, which was diluted with an equal volume of the same buffer before imaging. This procedure results in ALDs with monolayer mean molecular area (*MMA*) or area per PL values roughly comparable to cellular or physiological LDs (i.e., ~80 Å^2^/lipid, also referred to as *pMMA*) (48, 49), as quantified via lsFCS (see below). To obtain three more different *MMA* values, the PL to oil mass ratio was adjusted accordingly, e.g. 1/2000 PL/oil mass ratio for an increase in *MMA* as compared to *pMMA*, or 1/500 and 1/250 PL/oil mass ratios for a decrease in the *MMA* values. When needed, β-mercaptoethanol was mixed to this solution to a 5 % (v/v) final concentration for experiments in reducing environment. Next, AF488-labelled HCC was mixed to a final concentration of 200 nM and incubated for two hours. For Δ68, the final concentration was 500 nM. In the case of experiments conducted to compare the multimerization of Δ68 and HCC on the PL monolayer (i.e for experiments in which comparable binding to the ALD is needed for the two proteins), the latter was added to a final concentration of ~20 nM. Following the incubation step, the sample with the protein-ALD suspension was washed with 20 mM NaP to get rid of the unbound protein molecules. It is worth mentioning that cellular LD monolayers contain small amounts (0.4-1 mol%) of anionic PLs (50, 51). However, to model the HCV infection condition (in which anionic PLs are upregulated) and observe specifically the effect of specific anionic PLs on HCC binding, the model membranes studied in this work contained varying amounts of anionic PLs, up to 10 mol%.

### 5. Preparation of GUVs

GUVs as model of the ER bilayer (33) were used to investigate the binding behaviour of HCC to PLs. To highlight and compare the role of specific anionic PL, GUVs were prepared with the same PL compositions as ALDs by electroformation, using cylindrical Teflon chambers containing two platinum wires (52, 53) following an available protocol (39). PL solutions (2.5 mg/mL) were mixed with 0.005 molar% Rh-PE and 1 molar% biotinylated-PE. Following the electroformation, 100 µL of GUV suspension were transferred to an observation chamber, which was pre-treated with 100 µg/ml Streptavidin for 12 hours and then 10 % (w/v) BSA solution for 15 minutes. This suspension was further diluted with an equal volume of 40 mM NaP buffer and the GUVs were then allowed to settle down for 30 minutes, allowing the binding to the bottom of the plate. Next, AF488-labelled HCC was mixed to a final concentration of 200 nM and incubated for 2 hours. Finally, the GUVs were washed with 40 mM NaP to remove the unbound protein.

### 6. Line-Scan fluorescence correlation spectroscopy

Fluorescently labelled HCC binding, multimerization state and dynamics on ALDs and GUVs were quantified in terms of fluorescence intensity, brightness and diffusion coefficient (*D_Protein_*), respectively, using lsFCS. Furthermore, the same approach was used to measure PL diffusion coefficient (*D_PL_* or *D_PL/TF-PC_*) and packing density (indicated by the *MMA*) in ALD monolayers labelled with 0.001 molar% Rh-PE (or 0.005 molar% TF-PC).

LsFCS was performed on GUVs and LDs based on previous protocols (39, 54). Briefly, membranes were scanned perpendicularly with a pixel dwell time of 0.67 µs (pixel size 0.08 µm) at resolution of 256 pixels (total scan time ~4 min). Large-scale (ca. > 1µm) PL or protein clusters on the membrane were avoided in order to scan a relatively homogenous region. Importantly, unless a cluster was present, scanning was always performed at the lower right side of the membrane (135°) of each ALD or GUV.

All images were acquired with a Zeiss LSM880 system (Carl Zeiss, Oberkochen, Germany) using a Plan-Apochromat 40x/1.2 Korr DIC M27 water immersion objective and a 32-channel gallium arsenide phosphide (GaAsP) detector array. AF488-labelled HCC bound to ALD or GUV surface were imaged using a 488 nm Argon laser (0.77 µW at the exit of the objective). Analogous experiments in samples without proteins were performed to determine *MMA* and *D_PL_* in ALDs, using a 561 nm diode laser (1.97 µW). For measurements in the presence of HCC, *D_PL_* was determined simultaneously with the protein measurement on the same ALD, while for *D_PL/TF-PC_* determination, unlabelled HCC was used. For all measurements, the photobleaching and the background were kept below ca. 10 % / minute and 1% of the total initial intensity, respectively (54). In all cases, a commonly used bleaching correction and high-pass filters were applied (40).

In the case of ALD imaging, the significant difference in refractive index between the oil and aqueous phases results in imaging artifacts, such as enlargement of the point-spread function or unwanted reflections (causing e.g. the PL monolayers to appear as double lines, or a single abnormally broad line). Such distortions are stronger above the equatorial plane of ALDs partially adhering to the bottom glass coverslip. Imaging around the equatorial plane results in the best signal-to-noise ratio (48). Hence, all imaging was performed at this height. The validity of this approach was confirmed by comparing the PL packing density with that obtained using an alternative method described by Gandhi *et al*. (55), in which the ALDs are imaged at the south pole but they are not fixed to a coverslip (data not shown). Of note, such an approach could not be employed in general for this study, since the experiments described here require extensive washing to remove unbound proteins.

Prior to performing lsFCS, calibration of the excitation beam waist (*w_0_*) was performed each day by measuring the fluorescence autocorrelation curve of 0.25 µM AF488 in 20 mM NaP to correct the laser alignment variation. This calibration procedure was also performed for the PL (Rh-PE) experiments, using a similar concentration of Alexa Fluor^®^ 555 (Thermo Fisher Scientific, Waltham, MA, USA) (AF555) dye. Next, a mean of the diffusion times (*τ_Diff_*) from three independent autocorrelation curves was used to calculate *w_0_* by applying the reported diffusion coefficient (*D*) of the fluorescent dyes (56, 57) following **Eq. 1**:

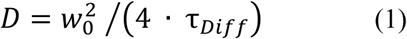

As an approximation, the ca. 2 % decrease in *D* due to the presence of the solute (58, 59) was neglected (39). Each data set was analysed as mentioned in previous work (39).

The effective illuminated area (*A_eff_*) on the GUV or ALD was estimated as (40):

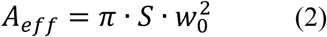

wherein the structural parameter is denoted by *S* (derived as mentioned in previous work (39)).

The analysis of each measurement resulted in the number of particles (*N*) in the measured area, the integrated line intensity (*I*) and ***τ****_Diff_* for the fluorophore (AF488 labelled HCC/ Rh-PE/ TF-PC). *I* was calculated as the time integral of the fluorescence signal in the immediate proximity of the membrane (60, 61) and expressed in Hz.

For the PL-only experiments, the total number of PL molecules in the illuminated area (*N_total_*) was determined, which was used to calculate the PL *MMA* on the ALD surface (39):

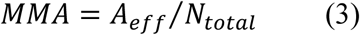

For the measurements with AF488 labelled HCC, the measured *I* was normalized through i) the brightness value of AF488 (*B_dye_*) obtained from daily calibration in solution (so to avoid errors caused by changes in laser power or optics alignment) and ii) the degree of labelling of the protein (so to correct for small sample-to-sample variations). In this regard, the typical *DOL* was ca. 0.2.

Protein brightness (*B_Protein_*) was determined as (62):

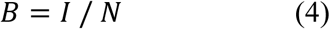

Hence, *B_Protein_* was expressed in Hz/molecule (Hz/mol) according to **Eq. 4**.

The *I* and *B_Protein_* values were converted into protein monomer concentration (mol/µm^2^) and multimerization state, respectively. In this context, the number of protein monomers in the detection area was calculated by dividing the respective *I* value by the monomer brightness (*B_monomer_*: in a rough approximation, the lowest *B_Protein_* value of all measurements). Next, the protomer concentration (i.e. protein concentration in units of equivalent monomer amounts) was obtained by dividing the number of protein molecules by the *A_eff_*. Finally, the total number of bound proteins was obtained by dividing the labelled protein concentration by the *DOL*. Hence:

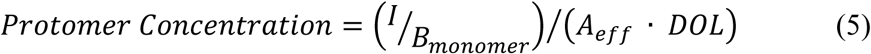

Following this, the multimerization state from each brightness value was calculated based on the reported equation (62):

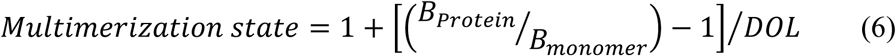

### 7. Transmission electron microscopy

ALDs (in Tris buffer) with or without HCC (200 nM) were negatively stained with uranyl acetate. Both samples were stained 2 hours after the preparation of the ALDs. 5 µL of sample were applied on a plain carbon grid, incubated for 45 seconds (s) and subsequently washed four times in 5 µL droplets containing 2 % (w/V) uranyl acetate dissolved in water. Afterwards, the sample was incubated for 30 s and the excess solution removed by blotting. For each sample more than 100 transmission electron microscope (TEM) micrographs were collected on a Talos F200C microscope (Thermo Fisher scientific, Waltham, MA, USA) with a CETA camera at a magnification of 22,000x using an exposure time of 1 s.

### 8. Software

Statistical significance of differences between data sets were calculated using one-way ANOVA or two-sample t-test (significance level=0.05). All figures and statistical tests were made using Origin Pro 2024b (OriginLab Corporation, Northampton, MA, USA) (38, 39). Fluorescence microscopy images were processed using Fiji software. All data were analysed with MATLAB 2020b (The MathWorks, Natick, MA, USA) using already available codes (39). Primer designing was performed using Benchling (San Francisco, California, USA).

## Results

### 1. Anionic PLs in LDs promote binding of HCC

HCV infection is associated to an increase in intracellular concentration of certain anionic PLs, such as PI(4)P and PI(4,5)P2 (8, 34, 36). However, it is not known whether these PLs play a role in HCC interactions with monolayers or bilayers. In order to understand the PL binding behaviour of HCC, we performed first a preliminary screening using immobilized PLs (i.e., PIP™ strips). PLs carrying different headgroups are presented on a nitrocellulose membrane and patterned as separate spots to test protein-PL interactions. Following this procedure, HCC binding was detected mostly within anionic PL spots with varying intensities, depending on the PL headgroup (**Figures 1A** and **S1**). The highest binding signal was observed for PI(4)P and PI(3)P, while that for the zwitterionic PL PC was among the weakest signals that could still be quantified (**Figure S1**). The results were reproducible, except for PI(3,4,5)P3, PI(3,4)P2, PI and PE which were, therefore, not further investigated. It is worth noting that the physical properties of PLs in this system, including their spatial organization and density, might differ from those of a PL monolayer on a water-oil interface and, therefore, must be accompanied by more in-depth investigation. For the following experiments, we thus focused on PI(4)P (as example of PIP with one phosphate group strongly interacting with HCC), PI(4,5)P2 (as example of PIP with two phosphate groups also interacting with HCC), PS (as an example of a non-PIP anionic PL still showing an interaction with HCC) and PC (as a negative control), as summarized in **Figure 1B**.

**Figure 1:**
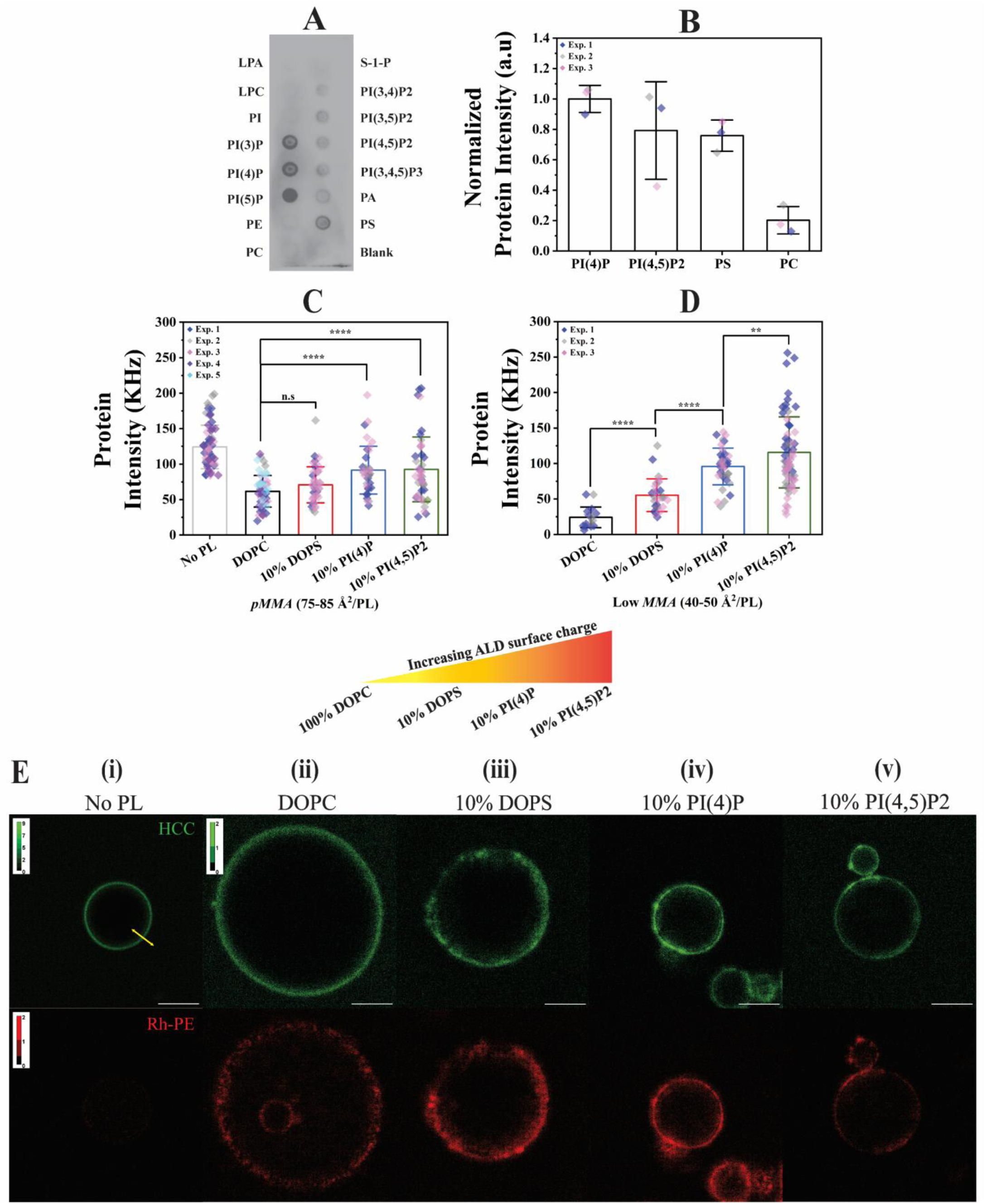
HCC binding to ALD monolayers depends on PL identity. **A** shows a typical binding pattern of HCC to different lipids in a PIP™ strip detected using complementary antibodies. **B** shows the quantification of HCC binding intensity to each spot from three independent experiments for four selected PLs. All data are normalized relative to PI(4)P. See **Table S1** for data points. **C** and **D** show the quantification of AF488-labelled HCC binding to ALDs performed via lsFCS and are represented as mean fluorescence intensity measured within the scanned region of the membrane (see Methods). **C** and **D** show the protein fluorescence intensity for ALD monolayers at *pMMA* and “low *MMA”* conditions, respectively. *MMA* values were not directly measured but rather estimated from protein-free ALDs prepared with the same PL/oil ratios (see **Table S6**). The ALDs are distinguished based on the interface PL composition: “No PL” refers to ALDs composed of only oil, “DOPC” to ALD monolayer composed of pure DOPC, “10% DOPS” to ALD monolayer composed of DOPC+DOPS (9:1 molar ratio), “10% PI(4)P” to ALD monolayer composed of DOPC+PI(4)P at 9:1 molar ratio and “10% PI(4,5)P2” to ALD monolayer composed of DOPC+PI(4,5)P2 at 9:1 molar ratio. All bars represent mean values with whiskers as the SD. Each point represents the result of one lsFCS measurement within one ALD. The points are pooled from at least two independent experiments denoted with individual colours. See **Tables S3**, **S4** and **S6** for details on statistics. Increasing surface charge trend of the ALDs as a function of different PL compositions is shown in form of a cartoon between panels **C**/**D** and **E**. For **C** and **D**, all statistical tests were performed via two-sample t-test at significance level 0.05 (**** p<0.0001, ** p<0.01, n.s: not significant). **E** shows representative fluorescence images of HCC-ALD interaction, in connection to the data in **C**. For each column, for the same samples shown in panel **C**, the green channel (top) represents AF488-labelled HCC, and the red channel (bottom) represents Rh-PE doped (0.001 molar%) ALDs. The yellow line in **(i)** depicts a typical scanning line used for lsFCS. The colour intensity range in the green channel for **(ii)**, **(iii)**, **(iv),** and **(v)** was rescaled with respect to that of **(i)** (lightest colour for pixels with 2 or more photons), for better visualization. Scale bar is 5 µm.

We proceeded to investigate HCC-lipid interactions directly in biophysical models of LDs by means of confocal microscopy and lsFCS. The latter measures the fluctuations in fluorescence emission from a spatially confined region of the membrane (see **Methods**) and calculates the labelled molecule mobility (in terms of *D* values), local abundance (in terms of the measured fluorescence signal *I* and the detected number of particles *N*) and multimerization state (in terms of brightness values) (40, 63). It is an advantage of this approach that brightness can be measured directly to determine the multimerization state, rather than relying on measurements of *D* that is, instead, only weakly dependent on protein mass (roughly, *D* ∝ Mass^−1/3^) (64).

First, we characterized protein-free ALDs to determine typical *MMA* values and the corresponding PL dynamics. By using multiple PL/oil mass ratios (see **Methods**), it is possible to cover an *MMA* range between ca. 40 and 120 Å^2^/PL (**Figure S2** and **Table S6**). It is worth noting that ALDs containing anionic PLs occasionally presented µm-scale lateral inhomogeneities (see e.g. **Figure S3A**) that were avoided during lsFCS measurements. PL mobility was quantified as a function of *MMA*, to confirm that the monolayer dynamics in our ALD model vary as expected (39, 65). Specifically, *D_PL_* values increased from ~5 µm^2^/s to ~9 µm^2^/s (~6 µm^2^/s for PIPs) for increasing *MMA* values, as expected for unsaturated PLs at the oil-water interface (66) (**Figure S2E**). Also, we could identify the best PL/oil ratios needed for each PL composition to obtain *MMA* values either in the range 75-85 Å^2^/PL, i.e. *pMMA* (48, 49) or in the range 40-50 Å^2^/PL (i.e., an arbitrarily chosen range, here defined as “low *MMA*”), as shown in **Table S6**. This estimation procedure is needed later for samples containing proteins, in which the *MMA* of each ALD cannot be directly measured (due to hindered PL dynamics and bleaching, see below). For example, to obtain *pMMA* samples, a PL/oil mass ratio of 1:1000 is required for DOPC ALDs, while 1:500 is more suitable for DOPC + PI(4)P (9:1) ALDs.

Next, we analysed HCC binding by quantifying the fluorescence signal of AF488-labelled HCC bound to PL monolayers formed at *pMMA*. For ALDs composed of the zwitterionic PL DOPC mixed with 10mol% of different anionic PLs, increasing surface charge led in general to increasing protein binding (**Figure 1C**). Here, we calculated that at pH 7.4, PI(4)P and PI(4,5)P2 possess a negative charge 2.94-fold and 4.2-fold higher than DOPS, respectively (67). Of note, control experiments with fluorescent PL analogues suggest that the different anionic PLs tested here partition into the monolayer in the expected amounts (**Figure S3**). A further control experiment indicates that, after washing the unbound protein away, the unbinding kinetics are significantly slower than the typical measuring time (**Figure S4**) and, therefore, the amount of bound protein can be reliably measured at “quasi”-equilibrium and is independent from the small amount of residual protein in solution.

Interestingly, the protein associates to the surface of ALDs even more efficiently if no PLs are present (“No PL” condition in **Figure 1**). For ALDs prepared from pure DOPC, “low *MMA*” values (i.e. higher PL density) are associated to a ca. 50% decrease in protein binding (cf. **Figures 1C** and **1D**, **Figure S5**), as expected from previous reports suggesting that PLs forms a barrier that prevents the interaction between protein and LD phase (33). Interestingly, we show that this effect was much less evident in the presence of anionic PLs i.e., ALDs containing 10 mol% DOPS (ca. 25% decrease in HCC binding) or PI(4)P (no decrease). Strikingly, **Figure S5** shows that protein binding even increased at higher PL packing density in ALD monolayers containing PI(4,5)P2, compared with the same ALDs at *pMMA*.

In order to clarify whether the observed interactions between HCC and anionic PLs are solely driven by non-specific electrostatic interactions, we prepared ALDs with similar charge densities (i.e. DOPC monolayers with 10mol% DOPS, 3.4mol% PI(4)P and 2.3mol% PI(4,5)P2). As shown in **Figure S6**, for p*MMA* samples (**Figure S6A**), PI(4,5)P2 appears to bind the protein slightly more efficiently than PS. The effect is more evident though in low *MMA* samples (**Figure S6B**), in which PI(4,5)P2 and, especially, PI(4)P exhibited a significantly higher affinity to HCC, compared to DOPS, despite similar charge density.

Finally, **Figure 1E** and **S6E** show typical confocal microscopy images of the examined ALDs at *pMMA*, in which the green channel indicates the AF488-labelled protein signal, and the red channel indicates the fluorescent PL signal (0.001 mol% Rh-PE). In line with lsFCS quantification (**Figures 1C** and **S6A**), HCC appears to bind efficiently to ALDs in the absence of PLs and much less when PLs form a monolayer at the surface of the ALD (note the different colour scaling in the images). A similar trend and a general agreement between confocal imaging and lsFCS is observed also at “low *MMA*” values/ high PL packing (**Figure S7**). Microscopy imaging additionally indicates that the lateral distribution of PLs and HCC is not homogeneous in the presence of anionic PLs.

### 2. PI(4)P promotes multimerization of HCC bound to LDs

Further analysis of HCC-ALD interactions via lsFCS indicates that the average protein multimerization (quantified via molecular brightness, see **Methods**) does not strongly depend on the tested PL compositions, at least at *pMMA* (**Figures S8** and **S9A**). It is worth mentioning that fluctuation-based quantification of protein multimerization was performed in regions of the PL membrane that did not contain µm-scale clusters (see previous paragraph), as this method is based on the analysis of the (fast) diffusion of fluorescent objects much smaller than the size of the optical point spread function (ca. 300 nm) (40, 63). At higher PL packing or “low *MMA*”, enhanced multimerization is observed for ALDs containing 10 mol% PI(4)P compared to e.g. 10 mol% DOPS or PI(4,5)P2 (**Figure S9B**) but, in general, a simple correlation between protein multimerization and *MMA* or PL identity is not evident over all the tested *MMA* values (**Figure S9C**). On the other hand, a clear trend is observed when plotting HCC multimerization as a function of local protein concentration (i.e., protein fluorescence intensity) (**Figure 2**). The observed molecular brightness values range from ~0.2 KHz/mol at low protein concentration to ~1 KHz/mol at the highest protein concentration measured, depending on PL composition. Considering that the protein labelling efficiency was ~0.2 dye molecules/protein and assuming that the lowest brightness we measured during this study (i.e. ~0.1 KHz/mol in GUVs, see below) corresponds to HCC monomers, it is possible to roughly estimate that HCC forms multimers with up to ca. 50 subunits in average, at concentrations ~1600 monomers/µm^2^ (**Figure S10A**) in the presence of PI(4)P. For DOPS and PI(4,5)P2 this number was 20-35 subunits in average. Interestingly, such concentration-dependent multimerization is less evident for pure DOPC ALDs. Next, an empirical model (68) was fit to the data in **Figure S10A** and the results confirm that the multimerization is the most efficient in ALDs containing PI(4)P, while no significant difference can be observed for ALDs containing DOPC, DOPS or PI(4,5)P2 (**Table S6 v**).

**Figure 2:**
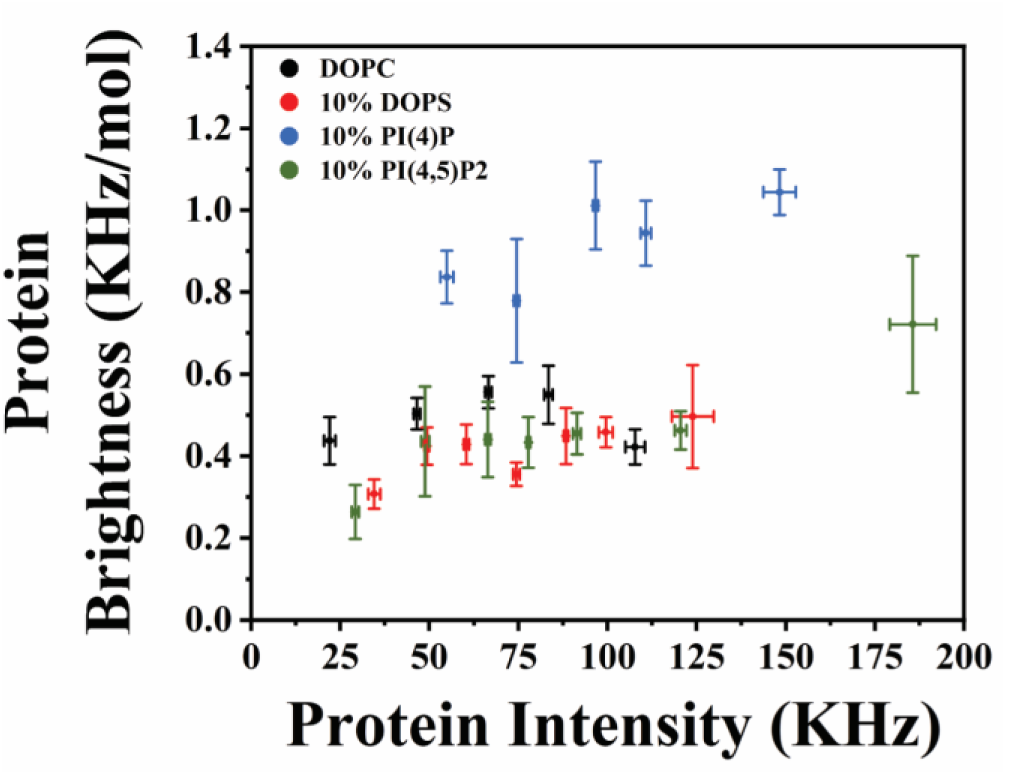
HCC brightness as a function of HCC concentration on ALDs. HCC multimerization is quantified via the protein brightness obtained via lsFCS analysis. The figure shows the protein brightness values for each ALD composition plotted as a function of the corresponding protein fluorescence intensity values. Each colour indicates the parameters measured on ALD monolayers made of DOPC, DOPC+DOPS (9:1), DOPC+PI(4)P (9:1) and DOPC+PI(4,5)P2 (9:1). For better visualization, the data points were binned so that each point represents the mean value calculated from 9-43 points for DOPC LDs, 10-28 points for 10% DOPS mixed ALDs, 7-30 points for 10% PI(4)P mixed ALDs and 13-37 points for 10% PI(4,5)P2 mixed ALDs. The error bars represent the SEM calculated from the binning procedure. See **Table S7** for the details on the data points.

Similarly to a previously published study (43), we aimed to test whether the protein-protein interactions observed here via lsFCS might be related to interactions which are relevant in the physiological context of virus capsid assembly. **Figure S11A** shows that, when bound to PI(4)P containing ALDs at similar concentrations as the wild-type protein, the assembly-incompetent HCC mutant Δ68 (44) exhibits a ca. 5- to 10-fold decreased brightness. Such brightness values are in the range of the lowest brightness values measured throughout this study (see below, for HCC bound to DOPC ALDs in reducing conditions, or HCC bound to PL bilayers) and might correspond to protein monomers or dimers. Also, for the samples in which we observed the most efficient protein multimerization (ALDs containing PI(4)P, as shown in **Figure S10A**), TEM imaging indicates the presence of regular structures with recurring spatial patterns with size between ca. 10 and 50 nm, that might correspond to the initial steps of viral capsid assembly (**Figure S12**).

Finally, we analysed the diffusive dynamics of HCC in ALDs under the investigated conditions (**Figures S10B** and **S13**). *D_Protein_* ranges from ~0.4 to ~1 µm^2^/s, with no strong correlation to protein concentration (**Figure S10B**), PL type (**Figures S13A** and **B**) or PL packing (**Figure S13C**).

In conclusion, the results so far indicate that negatively charged PLs (and, particularly, PIPs) promote HCC binding to ALDs. This effect is more easily observed if PLs are tightly packed within the monolayer. Furthermore, PI(4)P appears to enhance HCC concentration-dependent multimerization behaviour.

### 3. Cysteine bridges are needed for effective HCC binding to ALDs

It is known that the HCC forms dimers via inter-protein disulphide bonds (18). Such dimers have been shown to be important for capsid stabilization, virus particle production (5, 24) and HCC-membrane interaction (5). In comparison to non-reducing conditions (**Figures 1C** and **1E**), we found that the binding of HCC to ALDs, irrespective of the monolayer PL composition, is significantly hindered in a reducing environment, as evidenced by a ~5- to 10-fold decrease in the measured fluorescence intensity signal of the protein at the surface of ALDs (**Figures 3A** and **B**). Interestingly, lsFCS analysis reported a strong decrease in protein-protein interactions for DOPC ALDs (cf. **Figures S10A** and **3C**, **D**), as indicated by the low multimerization values measured in this sample. The multimerization values observed for ALDs containing anionic PLs are, on the other hand, comparable with those obtained before (**Figure S10A**, taking into account the lower concentration of the bound protein), including the fact that the largest multimers are observed in the presence of PI(4)P (e.g., between ca. 100 and 200 HCC protomers/µm^2^). Finally, the decrease in protein binding to ALDs does not appear to significantly modulate protein dynamics at the monolayer interface (cf. **Figures S13A** and **S14**).

**Figure 3:**
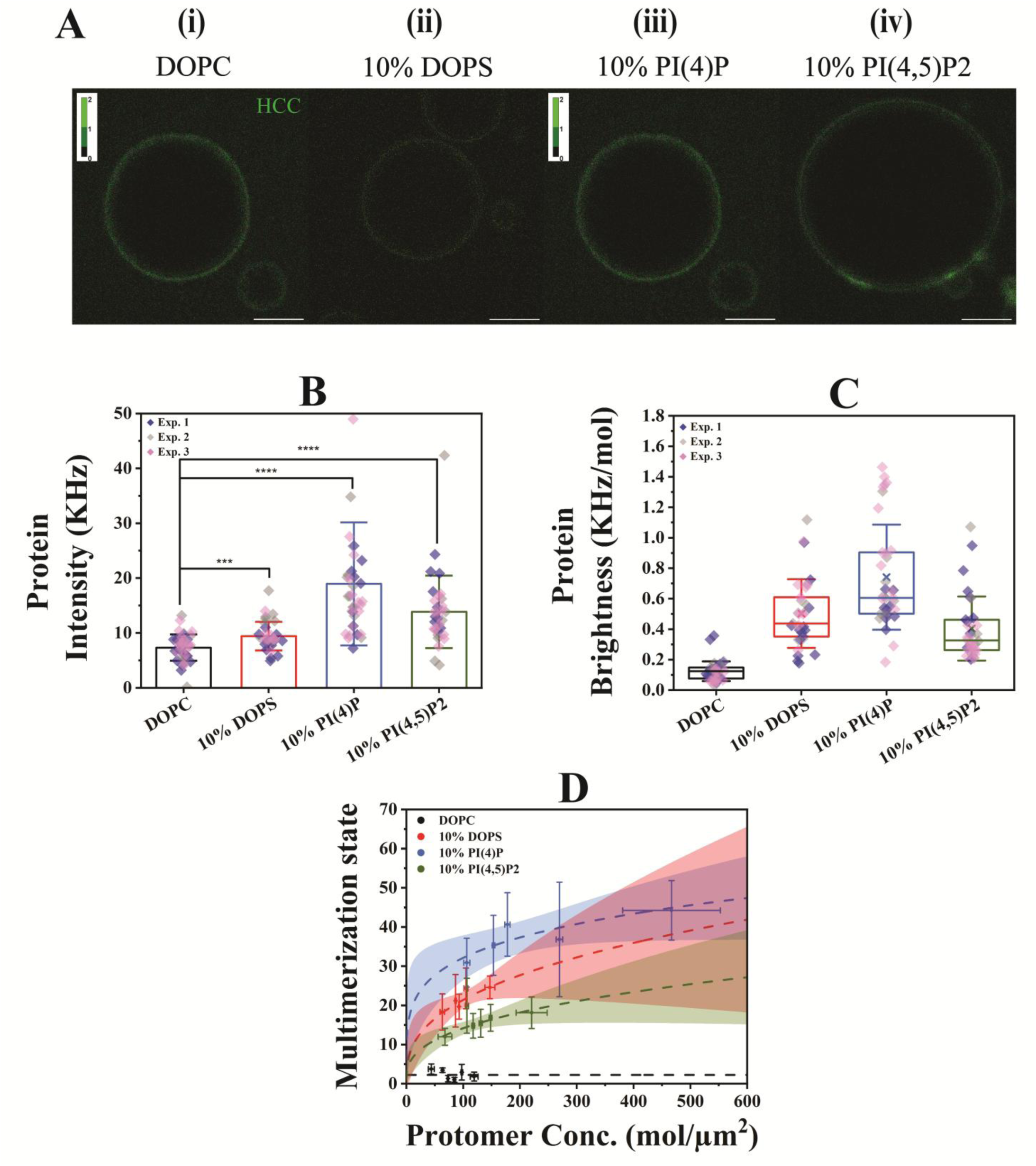
Decreased binding of HCC to ALD monolayers at *pMMA* in reducing environment. Quantification of AF488 labelled HCC binding and multimerization on ALD monolayers at *pMMA* condition was performed in reducing environment via lsFCS analysis and is represented as fluorescent protein intensity and brightness, respectively. **A** shows typical fluorescence images of HCC bound to ALDs in reducing environment, **(i)** for ALDs made from DOPC, **(ii)** for ALDs made from DOPC+DOPS (9:1), **(iii)** for ALDs made from DOPC+PI(4)P (9:1) and **(iv)** for ALDs made from DOPC+PI(4,5)P2 (9:1). The colour intensity range in the green channel for **(ii)**, **(iii)**, **(iv),** and **(v)** was rescaled with respect to that of Figure 1E **(i)** (lightest colour for pixels with 2 or more photons), for better visualization. The scale bar is 5 µm. **B** shows the fluorescent protein intensity on ALDs derived from images exemplified in **A**, as bar graphs. All bars represent mean values with whiskers as the SD. All statistical tests were performed via two-sample t-test at significance level 0.05 (**** p<0.0001 and *** p<0.001). **C** shows the protein brightness values of HCC after binding to the same set of ALDs, in the form of box plots. In all box plots, horizontal line is the median, ‘x’ marks the mean with first and third quartile as the boundaries and whiskers as SD. All data points were obtained from at least three independent experiments denoted with individual colours. **D** shows the protein multimerization state for each ALD composition plotted as a function of the corresponding protomer concentration values. Each colour indicates the parameters measured on ALD monolayers composed of DOPC, DOPC+DOPS (9:1), DOPC+PI(4)P (9:1), and DOPC+PI(4,5)P2 (9:1) respectively. For better visualization, the data points were binned so that each point represents the mean value calculated from 5-10 points for DOPC ALDs, 5-9 points for 10% DOPS mixed ALDs, 3-10 points for 10% PI(4)P mixed ALDs and 3-10 points for 10% PI(4,5)P2 mixed ALDs. Fitting was performed in the same way as for **Figure S10A**. 95% confidence interval range for each fit is shown by the respective colour bands. Protein concentration was obtained from protein intensity values and multimerization state was obtained from the corresponding brightness values using Eqs. 5 and 6 respectively. Error bars are SEM calculated from the binning procedure. The *a* value for 10% PI(4,5)P2 mixed ALDs is significantly lower than that for 10% DOPS and 10% PI(4)P mixed ALDs (p<0.05), as determined via one-way ANOVA at significance level 0.05. The *a* value difference between 10% DOPS and 10% PI(4)P mixed ALDs is not significant (p>0.05). Reducing environment was obtained through the addition of 5 % β-mercaptoethanol (v/v). See **Table S5** for the numerical values.

### 4. Anionic PLs are necessary for the binding of HCC to PL bilayers

In cells, removal of the SP from HCC results in its subsequent transfer from the ER bilayer to LD (25), which is the site of HCC multimerization - as a fundamental step of capsid assembly (15). Eventually, the HCC returns from the LD back to the ER bilayer (14, 26) and this re-targeting is crucial for efficient formation of virions (31). While it was reported that the D2 domain of HCC binds to bilayers exclusively in the presence of TG in the interleaflet region (33), the role of anionic PLs in this process has not been addressed yet. Therefore, we have investigated the interaction between HCC and GUVs containing different anionic PLs.

The binding of HCC to these bilayers is in general significantly (between one and two orders of magnitude) lower than to ALDs at *pMMA* (**Figures 4A** and **B**, cf. with **Figures 1E** and **C**). Interestingly, a fluorescence signal from the bound protein significantly above the background was observed exclusively in the presence of anionic PLs (**Figure 4A**). Probably due to the low concentration at the membrane, HCC does not significantly oligomerize, for all analysed compositions, as evidenced by multimerization states well below those measured for ALD-bound proteins (cf. **Figures S15** and **S10A**). LsFCS analysis also shows that diffusive dynamics of HCC on the GUV surface, i.e. *D_Protein_* values, are ~3 times faster than HCC bound to ALDs at *pMMA*, irrespective of the PL composition (**Figure 4C**). It is worth noting that HCC brightness (or multimerization) measurements and *D* quantification were not possible for GUVs containing exclusively DOPC, as the fluorescence signal from the bound protein was not sufficiently high for lsFCS analysis.

**Figure 4:**
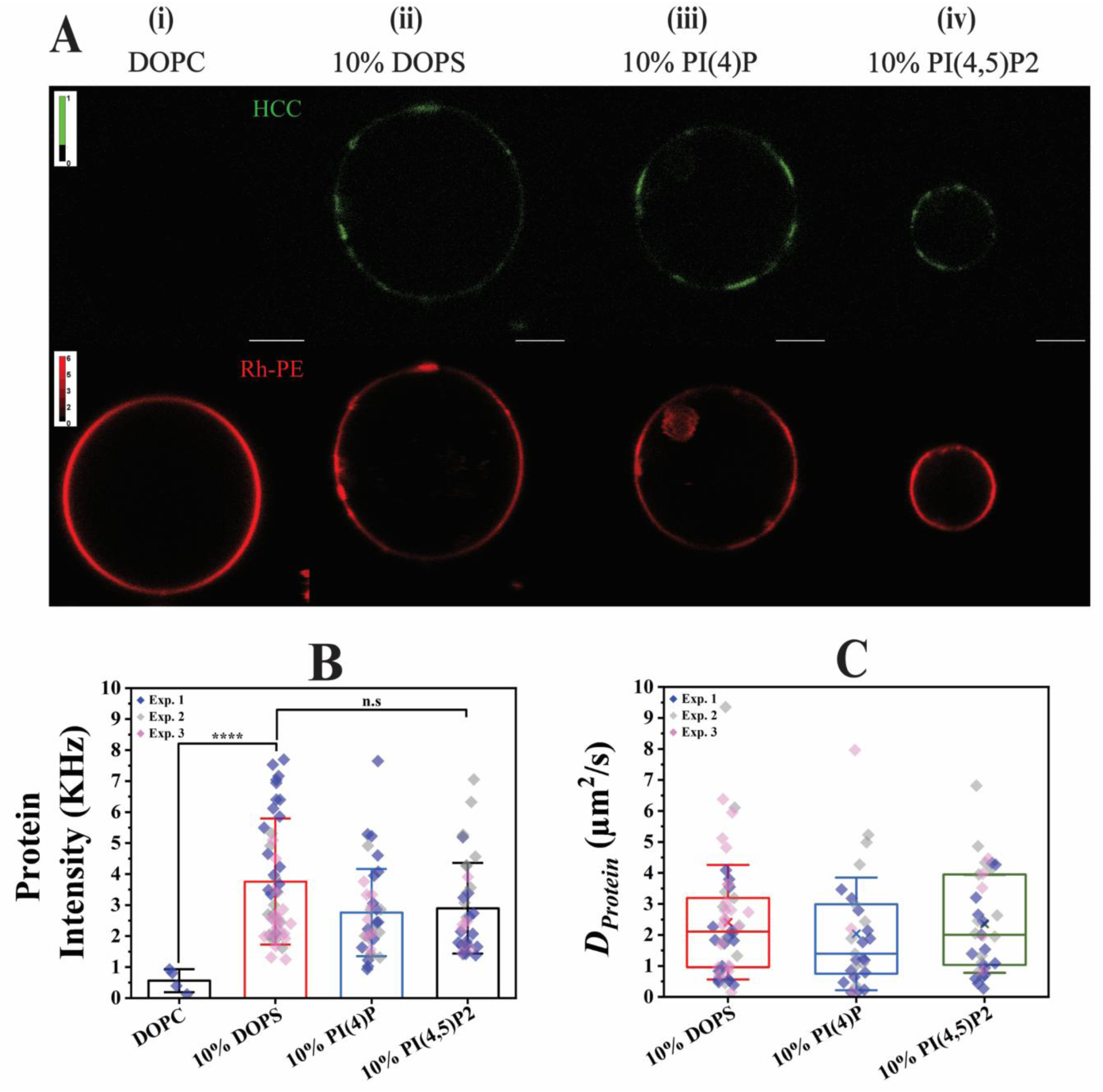
HCC binds to bilayer models only in the presence of anionic PLs. Quantification of AF488 labelled HCC binding, multimerization and mobility via lsFCS analysis on GUVs composed of different PL species represented as protein intensity and diffusion coefficient (*D_Protein_*). **A** shows typical fluorescence images of HCC bound to GUVs: **(i)** for GUVs made from DOPC, **(ii)** for GUVs made from DOPC+DOPS (9:1), **(iii)** for GUVs made from DOPC+PI(4)P (9:1) and **(iv)** for GUVs made from DOPC+PI(4,5)P2 (9:1). For each column, the green channel (top) shows the AF488 labelled HCC, and the red channel (bottom) shows Rh-PE labelled GUVs. The colour intensity range in the green channel was rescaled with respect to that of Figure 1E **(i)** (lightest colour for pixels with 1 or more photons), for better visualization. The red channel was also rescaled compared to Figure 1E **(i)** (lightest colour for pixels with 6 or more photons). Scale bar is 5 µm. **B** shows the fluorescent protein intensity on GUVs quantified from images exemplified in **A**, shown as bar graph. All bars represent mean values with whiskers as the SD. **C** shows the distribution of *D_Protein_* values from HCC binding to the same set of GUVs (except DOPC) in the form of box plots. GUVs of all PL compositions were labelled with 0.005 molar% Rh-PE for visualization purpose. In all box plots, the horizontal line is the median, ‘x’ marks the mean with first and third quartile as the boundaries and whiskers as SD. All data points were obtained from three independent experiments denoted with individual colours. In **B**, the differences between 10% DOPS, 10% PI(4)P and 10% PI(4,5)P2 mixed GUVs are statistically not significant. In **C**, mean *D_Protein_* values are not distinguishable. All statistical tests were performed via two-sample t-test at significance level 0.05 (**** p<0.0001 and n.s: not significant). See **Table S8** for the numerical values.

### 5. HCC binding hinders PL dynamics in ALD monolayers

In order to test the effect of protein binding on PL dynamics, we monitored the diffusion of the fluorescent PL analogue Rh-PE (*D_PL_*). The underlying assumption here is that the viscosity and the local membrane environment sensed by the fluorescent PL are a good approximation of those sensed by the rest of the PLs (39). Irrespective of the PL composition, *D_PL_* in ALD monolayers at *pMMA* is strongly decreased (~10-fold) following HCC binding (**Figure 5, Figure S16**). Interestingly, a decrease in lateral dynamics was also observed for a structurally different fluorescent PL analogue TF-PC (*D_PL/TF-PC_*), albeit by a much smaller extent (i.e., ~3-fold decrease) (**Figure S16C**).

**Figure 5:**
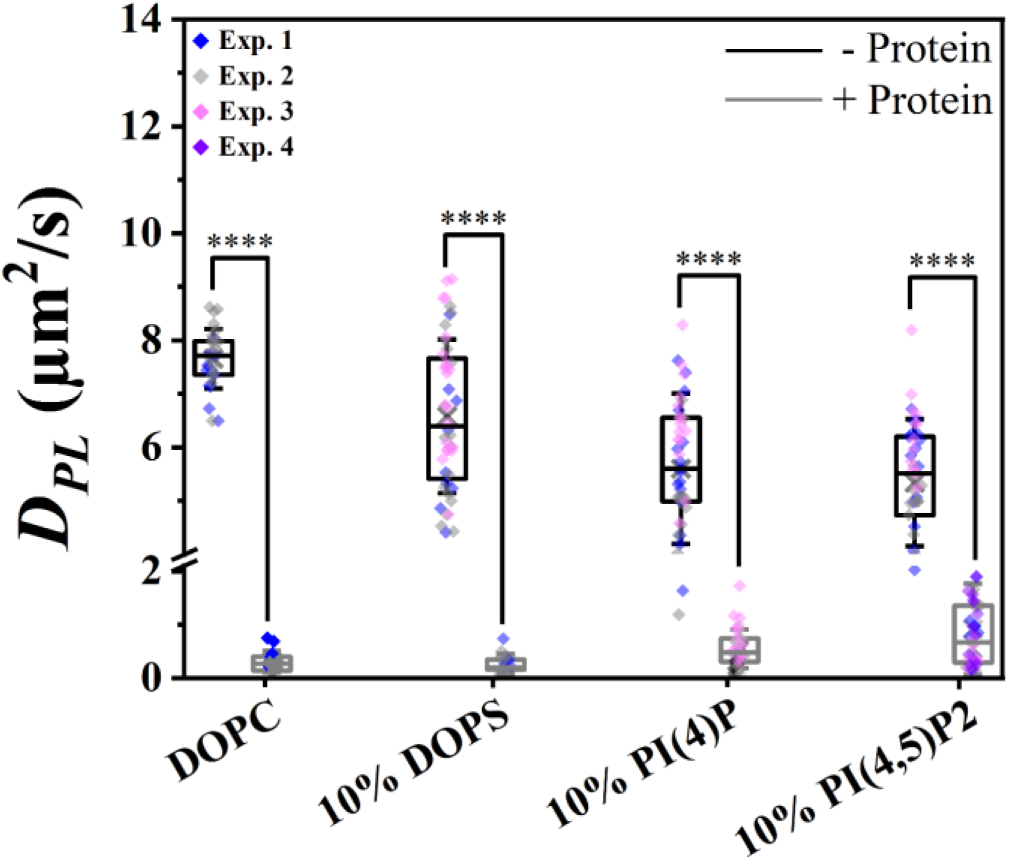
ALD monolayer PL mobility decreases as a consequence of HCC binding. PL mobility in ALD monolayers composed of DOPC, DOPC+DOPS (9:1), DOPC+PI(4)P (9:1) and DOPC+PI(4,5)P2 (9:1) at *pMMA* (labelled with 0.001 molar% Rh-PE), represented as *D_PL_* was analysed by lsFCS. All values are shown in the form of box plots with the data points from two to three independent experiments. The data points for ALDs without HCC are depicted in black boxes and those in the presence of HCC in grey boxes. In all box plots, the horizontal line marks the median, ‘x’ marks the mean with first and third quartile as the boundaries and whiskers as SD. Statistically different mean *D_PL_* values are shown with ****. All statistical tests were performed via two-sample t-test at significance level 0.05 (**** p<0.0001). See **Table S10** for the numerical values.

## Discussion

The binding of the HCC protein to the LD monolayer surface and subsequent return to the ER bilayer membrane is a fundamental step of the HCV capsid assembly (4, 14, 15, 26). In this complex process, the role of lipid molecules is not known yet, despite the evidence of significant upregulation of certain PL species in the HCV infectious cycle (8, 34, 69). In this work, we aimed to investigate the possible function of specific lipids in HCC-membrane interaction and subsequent assembly, as previous studies regarding HCC-LD interaction did not address this aspect (16, 33, 70, 71). A more recent investigation by Ajjaji and colleagues (33) highlighted the importance of triglycerides in the interaction between HCC and monolayers or bilayers. However, these results were based on the investigation of ALD monolayers composed of only DOPC and DOPE, thereby not addressing the possible effect of other (e.g. anionic) PLs, which are synthetized more abundantly during HCV infection (36). In this context, it is worth noting that measuring the precise amount of transient and precarious PLs such as PIPs might be quite complex (72). Furthermore, local concentration of PIPs might be dramatically increased by preferential partitioning into lipid domains (37, 73–75). Therefore, we have explored in this work a relatively large anionic PL concentration range (2-10 mol%), in general higher that previous estimates of average relative concentrations (0.4-1 mol% (50, 51)), so to obtain a complete picture of possible lipid-protein interactions.

Our experiments suggest that, irrespective of PL composition, the LD monolayer constitutes in general a passive barrier to HCC-LD interaction, as HCC shows the highest binding to oil droplets devoid of PLs. In analogy, HCC binding decreases dramatically, as more and more DOPC molecules are included in the monolayer (**Figure S5**). A previous study showed a similar effect, albeit for a specific domain (i.e., D2) of this protein (33). The oil-protein interaction seems therefore to dominate over possible PL-protein interactions (**Figure 1**). As monolayer might have packing defects (76), HCC might use such defects to interact with the oil phase (possibly via its amphipathic helix), rather than with (anionic) PLs (15, 33). Similar behaviour is observed for several LD-binding proteins (76, 77). Nevertheless, even at *pMMA* conditions, HCC binding to the ALD correlates with surface charge, assuming that each anionic PL contributes a negative charge increasing in the order DOPS<PI(4)P<PI(4,5)P2. In agreement with this hypothesis, HCC interactions with anionic PLs should be easier to detect if the interactions between protein and oil phase are hindered by a tighter packing in the monolayer. Indeed, increasing monolayer PL density enhances HCC-LD binding in the presence of e.g. PI(4,5)P2, opposite to what was observed for the zwitterionic DOPC (**Figure S5** and **1D**). Also, HCC affinity towards anionic PLs does not appear to be driven exclusively by electrostatic interactions, as we detected a stronger binding of the protein to PI(4)P-containing monolayers, among samples with comparable negative surface charge (**Figure S6B)**. PI(4)P appears to play a specific role also in modulating protein-protein interactions, as indicated by the enhanced concentration-dependant multimerization of HCC in ALDs containing this PIP, compared to other anionic PLs possessing lower (i.e., DOPS) or higher (i.e., PI(4,5)P2) negative charge (**Figure 2** and **S10A**). Of interest, in our ALD models, HCC forms multimers of up to ca. 50-60 subunits in average (at a concentration of ca. 2500 mol/µm^2^ and in the presence of PI(4)P), i.e. with a size similar to what is expected for the forming HCV capsid (44, 78–80). The apparent role of PI(4)P in facilitating capsid formation might relate to the observation that HCV infection is marked by high intracellular concentrations of this PL (36).

The observation that anionic PLs in the LDs (and, specifically, PI(4)P) might increase HCC-HCC interactions is further strengthened by experiments in reducing conditions (**Figure 3**). While inter-HCC cysteine bridges have been proposed to stabilize the capsid and thereby control the viral assembly process (5, 24), our data further suggest a role in promoting protein binding to the monolayer. Also, while protein multimerization is, as expected, strongly decreased in reducing conditions for the case of DOPC monolayers, HCC multimers remain instead apparently stable in the presence of anionic PLs, especially PI(4)P (**Figure 3D**). Once more, these observations suggest that HCC-HCC and HCC-LD interactions are enhanced by anionic PLs, albeit not exclusively due to non-specific electrostatic interactions.

In the context of HCC multimerization on the surface of ALDs, as detected via fluorescence fluctuation analysis, it is important to determine whether the observed protein-protein interactions are related to non-specific random protein aggregation or, rather, to physiologically relevant interactions. The latter possibility is supported by our control experiments. First, we have shown that the multimerization detected via lsFCS for the assembly-incompetent mutant HCC Δ68 is, as expected, strongly decreased (44, 81) (**Figure S11**). Second, it would not be easy to reconciliate the hypothesis of random protein aggregation (e.g. at high protein density) with the observation of higher multimerization specifically in the presence of PI(4)P (instead of e.g. PI(4,5)P2). Third, TEM imaging indicates the presence of protein structures with regular spatial features, e.g. in the shape of crescents, that are compatible with the hypothesis of ordered protein-protein interactions, rather than non-specific protein aggregation (**Figure S12**). We propose therefore that the multimers reported in this work might represent indeed physiologically relevant HCC-HCC interactions and, possibly, correspond to the initial stages of HCV capsid formation. While fluorescence-based imaging approaches with higher spatial resolution (e.g. STED) (82, 83) might help further investigating this issue, it must be noted that the low label density (i.e. *DOL*) used in this work would represent a significant problem for such approaches (84). Also, a higher label density might hinder or alter the expected protein-protein interactions, as reported e.g. for the HIV-1 capsid (85).

Regarding the molecular mechanisms driving protein-LD interaction, the observations of HCC binding to tightly packed monolayers and the high affinity of the protein for the oil phase of ALD devoid of PLs are compatible with the hypothesis that the protein interacts with the oil phase and, concurrently, to anionic PLs via specific electrostatic interactions. Also, the presence of direct, albeit weak, interactions between HCC and anionic PLs is confirmed by our experiments on model bilayers (**Figure 4**). While triglycerides were shown to promote the binding of HCC to bilayers (33) (e.g. to the ER), we demonstrate here that the presence of small concentrations of anionic PLs is also sufficient to obtain a protein binding significantly higher than that observed in the presence of only zwitterionic PLs. Of note, HCC binding to bilayers remains nevertheless much lower than the one observed in the case of monolayers and, as expected, is associated to much weaker protein-protein interactions (due to lower local concentration on the membrane, see **Figure S15**). In this context, the much slower diffusive dynamics of HCC on monolayers compared to bilayers (cf. **Figure 4C** and **S13**) suggest that the protein binds only superficially to bilayers (i.e. to PL headgroups (39)). On the other hand, in monolayers, the protein might at least partially insert into the highly viscous oil-phase (33), thus slowing down its lateral diffusion.

The last part of our work focuses on the effect of protein binding on the properties of the PLs in the LD monolayer. Since we hypothesize that direct interactions might exist between anionic PLs and oligomers (or large multimers) of HCC, it is reasonable to expect that protein binding might affect the lateral diffusion of PLs in the membrane (86, 87). Furthermore, it was also reported that PL dynamics is affected by the insertion of proteins into a monolayer or a membrane leaflet (87, 88). Accordingly, our results indicate a strong decrease in diffusive dynamics of a fluorescent PL probe (Rh-PE) after binding of HCC to the monolayer (**Figure 5**). Interestingly, the diffusion of a structurally different fluorescent PL (TF-PC) also decreases in the presence of a protein, but by a much smaller extent. Assuming that Rh-PE carries a negative charge at pH 7.4, while TF-PC remains neutral, the latter PL probe would report a general decrease in diffusive dynamics for all PLs in the monolayer (due e.g. to the insertion of the protein and the consequent increase in PL packing). On the other hand, the dramatic decrease of diffusive dynamics of Rh-PE might instead indicate that this anionic fluorescent PL (similarly to the other anionic PLs in the monolayer) binds to the bulky protein multimers and, as a consequence, diffuses at a much slower rate.

## Conclusions

This work shows that anionic PLs, specifically PI(4)P, affect HCC multimerization on monolayers (as models of LDs), probably by modulating the local concentration of the protein. In this scenario, HCC can be hypothesized to interact with the neutral lipid phase of LD and concurrently, although less strongly, with the polar heads of anionic PLs. Such significant involvement of anionic PLs could be one of the reasons why these are more abundant during HCV infection (8, 34, 36). Furthermore, we show that, through insertion in the monolayer or by recruiting specific PLs, HCC has the potential to alter the biophysical properties of LD monolayers. While the simplified model system used in this work was necessary to pinpoint specific lipid-protein interactions, future studies should investigate the role of more complex, physiologically relevant PL compositions and, most importantly, the role of other host or viral proteins.

## Author contributions

TM and SC conceived the research. TM, BA, SA, JR, PW and PJ performed the experiments and analysed the results. SC wrote the MATLAB codes for data analysis. TM and SC wrote the manuscript.

## Supporting information

Raw data

## Acknowledgements

TM would like to thank the Postgraduate Scholarship Committee of the University of Potsdam, Germany for his Doctoral dissertation completion scholarship and funding this project. We thank the members of the group, especially Dr. Anja Thalhammer for their valuable comments and discussions. Additionally, TM would like to thank Nadine Brandt for her help in preparing the supplemental tables and Melanie Anding for help with the TEM experiments.

## Declaration of interests

The authors declare no competing interests.

## Declaration of generative AI and AI-assisted technologies in the writing process

During the preparation of this work, TM used GPT.UP (OpenAI GPT-40) in order to improve the readability and language of the abstract and significance. After using this tool/service, the authors reviewed and edited the content as needed and take full responsibility for the content of the publication.

## Notes

### Competing Interest Statement

The authors have declared no competing interest.

### Summary of Updates

We have performed some more experiments to validate our results. This involves protein binding comparison on ALDs with similar charge. Additionally, we have also added TEM images of ALDs and HCC.

