## Supplemental file for "Phosphatidylinositol Phospholipids Modulate Hepatitis C Virus Core Protein Assembly on Lipid Membranes"

### Supplemental Figures

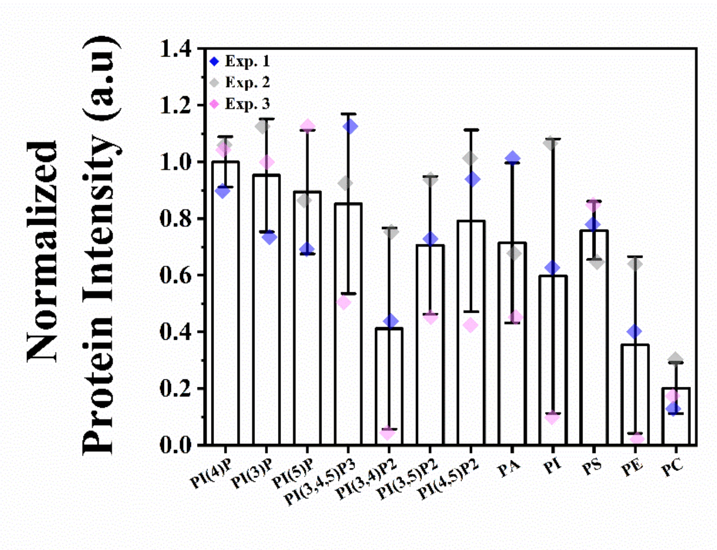


**Figure S1: Normalized HCC binding intensity within lipid spots of PIP™ strip.**
HCC binding to each PL spots on the PIP™ strip (**Figure 1A**) shown after normalization to the PI(4)P value. Protein binding signal was obtained from the chromogenic detection of a complementary antibody (see **Methods**). Bars represent mean values with whiskers as the SD. Data points were obtained from three independent experiments denoted with individual colours. See **Table S1** for details on statistics.

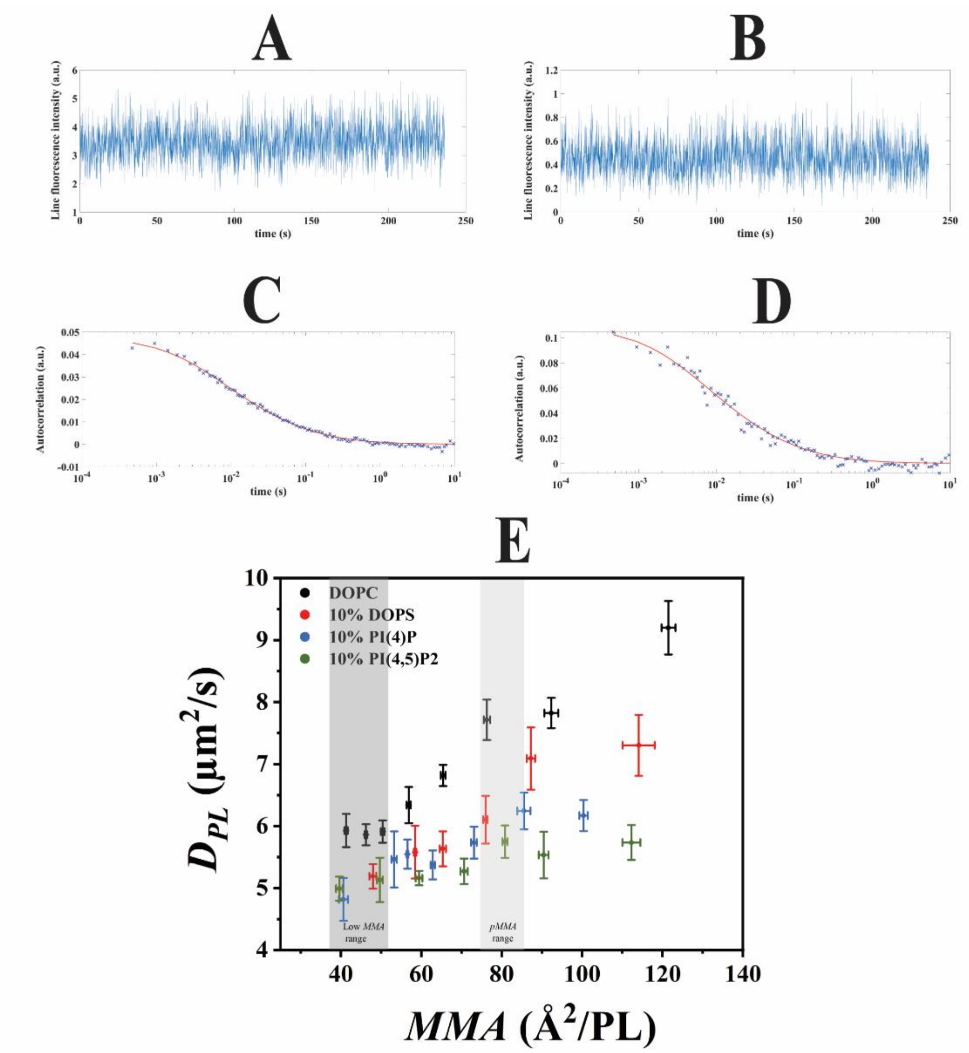


**Figure S2: PL mobility in protein-free ALDs increases as a function of decreasing packing density.**
**A** and **B** are representative fluorescence intensity vs. time plots of lsFCS measurement of Rh-PE molecules in DOPC and DOPC+PI(4)P ALDs, respectively, at *pMMA* condition. **C** and **D** show the corresponding autocorrelation curves, including a fit (solid line) to a 2D diffusion model. **E** shows PL mobility and packing density, represented as $D_{PL}$ and *MMA* respectively, and obtained via lsFCS analysis (1). All monolayers were doped with 0.001 molar% Rh-PE. $D_{PL}$ is plotted as a function of *MMA*. For each PL, ALD monolayers with different estimated *MMA*s were produced by adjusting the PL/oil ratio (see **Methods**). For better visualization, data points were binned so that each point represents the mean value calculated from 13-29 experimental points for DOPC ALDs, 12-37 points for 10% DOPS-containing ALDs, 9-19 points for 10% PI(4)P-containing ALDs and 13-20 points for 10% PI(4,5)P2-containing ALDs. Light grey and dark grey boxes show the *pMMA* range and "low *MMA*" range respectively. Each colour represents the parameters obtained from ALD monolayers formed of DOPC, DOPC+DOPS (9:1), DOPC+PI(4)P (9:1) and DOPC+PI(4,5)P2 (9:1). Error bars are SEM. See **Table S6** for data points.

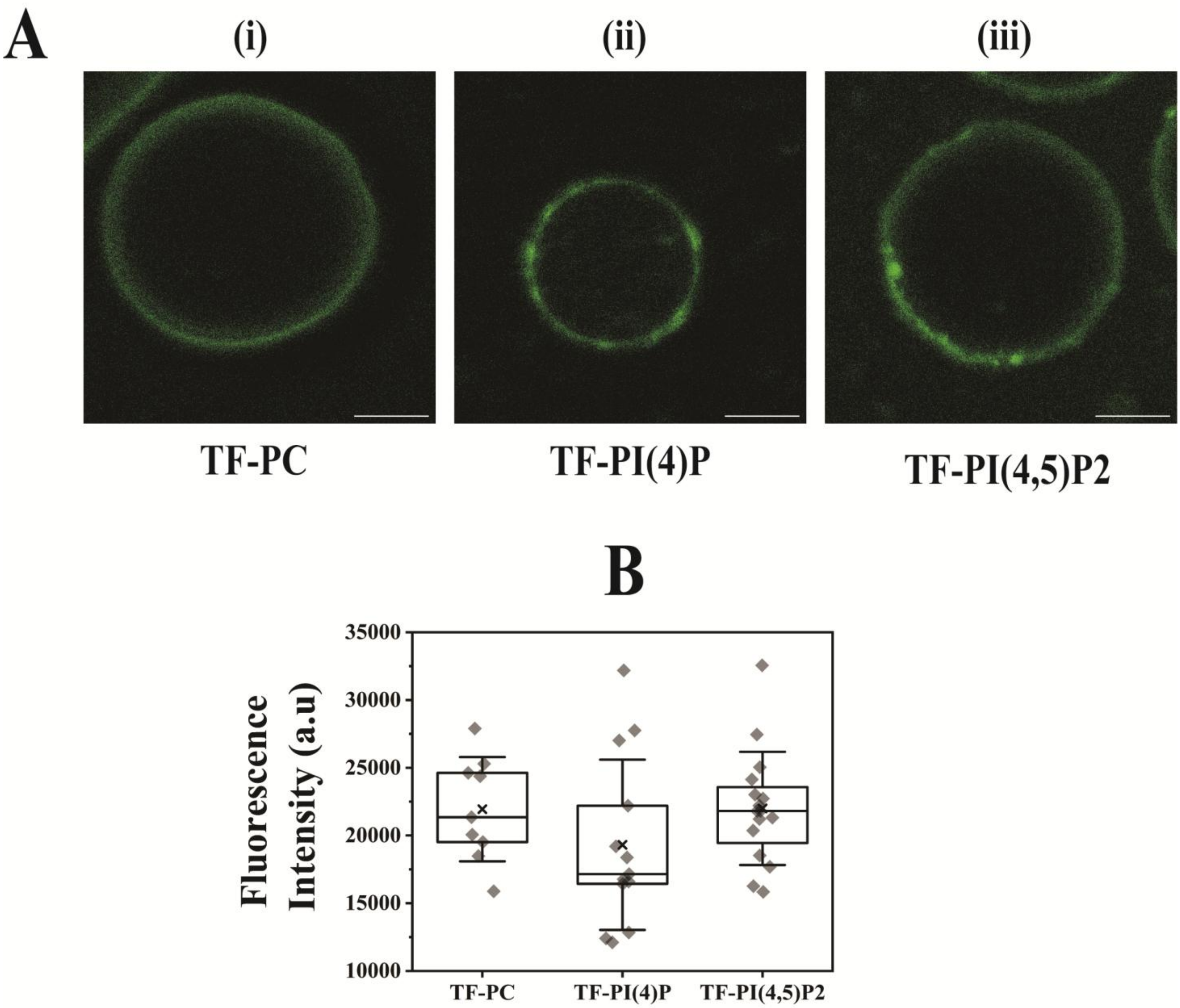


**Figure S3: Inclusion of different (charged) PLs into LDs is not strongly affected by PL identity.**
**A** shows typical fluorescence images of ALD monolayers under *pMMA* conditions labelled with TopFluor® -PC (TF-PC) **(i)**, TF-PI(4)P **(ii)** and TF-PI(4,5)P2 **(iii)**. All monolayers were composed of DOPC and 0.005 molar% of the respective fluorophore. The colour intensity range was reduced by a factor of 4.5 with respect to that of **Figure 1E (i)** (green channel) using Fiji software, for better visualization. Scale bar is 5 µm. **B** shows the distribution of monolayer fluorescence intensity values of at least 9 ALDs of each fluorophore type i.e. TF-PC, TF-PI(4)P or TF-PI(4,5)P2 represented in the form of box plots. Fluorescence intensities were quantified using Fiji software (2) and MATLAB. In all box plots, the horizontal line represents the median value, while the 'x' indicates the mean with the first and third quartile as the boundaries. The whiskers indicate SDs. Both zwitterionic and anionic TopFluor® derivatives partition in the ALD monolayer as expected, as shown by the similar fluorescence intensities in **B** (determined via one-way ANOVA, p= not significant at significance level 0.05). See **Table S6(i)** for the data points.

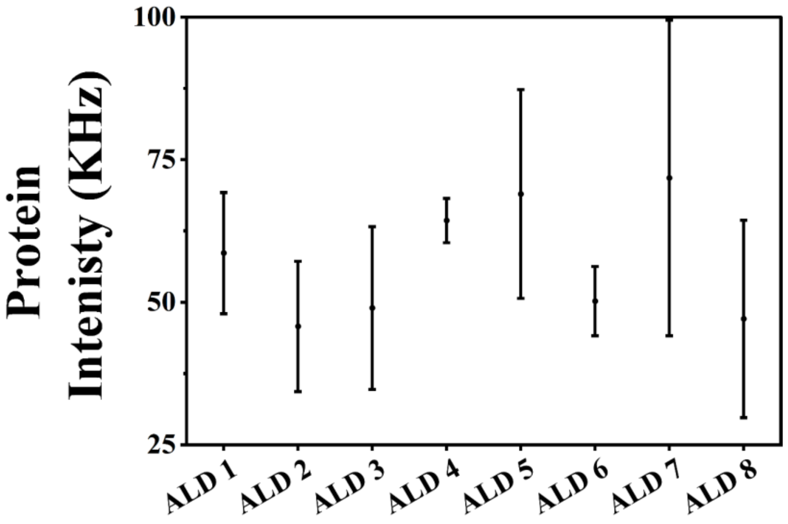


**Figure S4: Variation of initial fluorescently labelled protein intensity during the course of a typical measurement.** This figure shows, as an example, the initial fluorescence intensity values of HCC bound to DOPS-containing ALDs at low *MMA*, after removal of the unbound protein (data from **Figure1D** 10% DOPS bar). This specific sample was chosen because we observed the lowest affinity between DOPS and HCC, among all measured anionic PLs. The value for each ALD, acquired at the beginning of each lsFCS measurement, is shown as a function of the measurement order. Roughly, a time gap of ca. 5 minutes existed between each measurement: ca. 4 minutes for a lsFCS measurement plus 1 minute to locate the next ALD. If fast significant protein dissociation took place during the whole measurement pipeline, we would expect a negative correlation between bound protein signal and number of analysed ALD. On the contrary, no strong intensity decrease is observed over ca. 40 minutes. Each point is the mean of protein intensities measured for ALDs from three independent experiment days. Error bars are SEM. See **Table S2** for data point details.

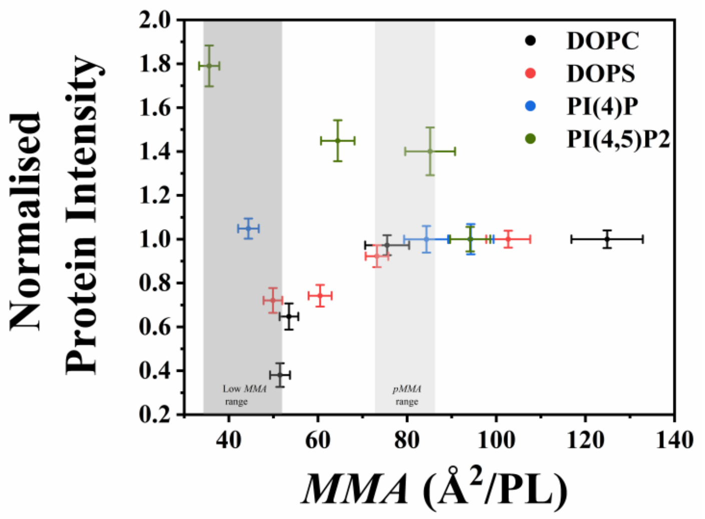


**Figure S5: Interplay between HCC binding and monolayer *MMA* is determined by PL composition.** Quantification of AF488-labelled HCC binding to LDs composed of DOPC, DOPC+DOPS (9:1), DOPC+PI(4)P (9:1) and DOPC+PI(4,5)P2 (9:1) was performed via lsFCS and represented as fluorescent protein intensity. The normalized values are represented as a function of ALD monolayer *MMA*, which, in turn, were determined for 4 PL/oil ratios and measured via lsFCS (see **Methods** and **Table S5**). Here, each point represents the mean values of each of the two parameters obtained from at least 18-59 experimental points for DOPC ALDs, 28-51 points for DOPS mixed ALDs, 37-40 points for PI(4)P mixed ALDs and 38-69 points for PI(4,5)P2 mixed ALDs with error bars as SEM. *MMA* values were not directly measured but rather estimated from protein-free ALDs prepared with the same PL/oil ratios (see **Table S5**). Light grey and dark grey boxes show the *pMMA* range and "low *MMA*" range respectively. All monolayers were labelled with 0.001 molar% Rh-PE. See **Table S6** for details on the statistics.

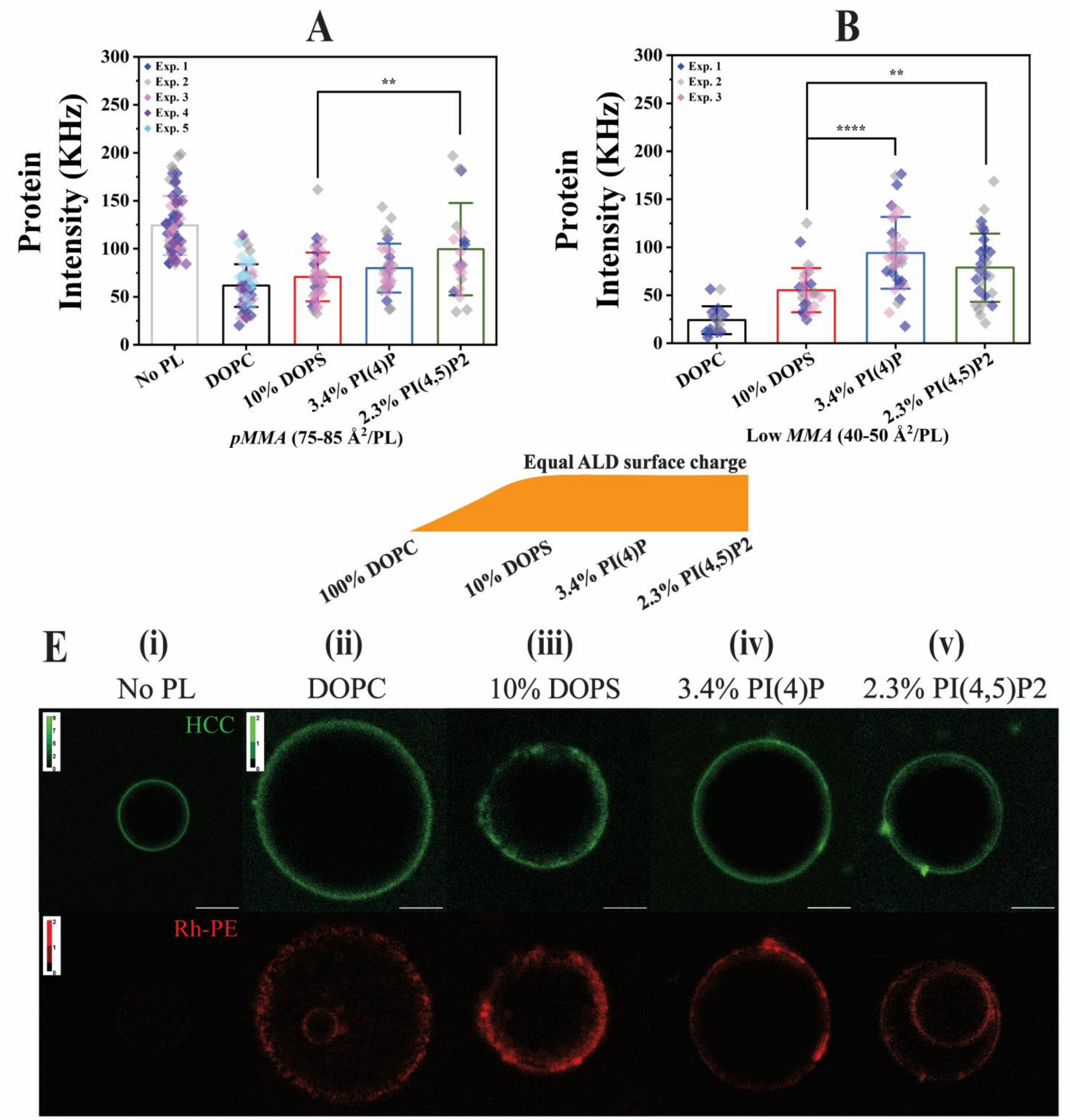


**Figure S6: HCC binding to ALD monolayers carrying comparable surface charge.**
**A** and **B** show the quantification of AF488-labelled HCC binding to ALDs performed via lsFCS and are represented as mean fluorescence intensity measured within the scanned region of the membrane (see Methods). **A** and **B** show the protein fluorescence intensity for ALD monolayers at *pMMA* and "low *MMA"* conditions, respectively. *MMA* values were not directly measured but rather estimated from protein-free ALDs prepared with the same PL/oil ratios and (see **Table S5**). The LDs are distinguished based on the interface PL composition: "No PL" refers to ALDs composed of only oil, "DOPC" to ALD monolayer composed of pure DOPC, "DOPS" to ALD monolayer composed of DOPC+DOPS (9:1 molar ratio), "3.4% PI(4)P" to ALD monolayer composed of DOPC+PI(4)P at 9.66:0.34 molar ratio and "2.3% PI(4,5)P2" to ALD monolayer composed of DOPC+PI(4,5)P2 at 9.77:0.23 molar ratio. ALDs mixed with 10% DOPS, 3.4% PI(4)P and 2.3% PI(4,5)P2 carry approximatively the same surface charge, as shown by the cartoon between panels A/B and C. All bars represent mean values with whiskers as the SD. Each point represents the result of one lsFCS measurement within one ALD. The points are pooled from at least two independent experiments denoted with individual colours. See **Tables S1**, **S2** and **S3** for details on statistics. All statistical tests were performed via two-

sample t-test at significance level 0.05 (**** p<0.0001 and ** p<0.01). **C** shows representative fluorescence images of HCC-ALD interaction, in connection to the data in **A**. For each column, for the same samples shown in panel **C**, the green channel (top) represents AF488-labelled HCC, and the red channel (bottom) represents Rh-PE doped (0.001 molar%) ALDs. The colour intensity range in the green channel for **(ii)**, **(iii)**, **(iv),** and **(v)** was rescaled with respect to that of **(i)** (lightest colour for pixels with 2 or more photons), for better visualization. Scale bar is 5 µm. Data from "No PL", "DOPC" and "DOPS" are same as from **Figure 1**.

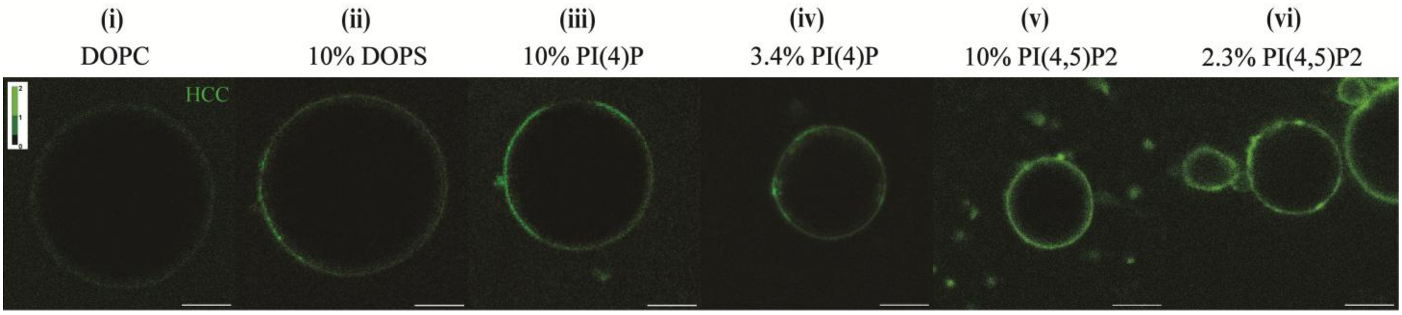


**Figure S7: HCC binding to ALD monolayers with high PL packing depends on PL identity.**
**(i)-(vi)** show representative fluorescence images of AF488-labelled HCC binding to ALD monolayers with "low *MMA"* (i.e., 40-50 Å²/PL), composed of DOPC, DOPC+DOPS (9:1), DOPC+PI(4)P (9:1), DOPC+PI(4)P (9.66:0.34) DOPC+PI(4,5)P2 (9.77:0.23) and DOPC+PI(4,5)P2 (9:1) respectively. These images exemplify the results shown in **Figure 1D** and **S6B**. The colour intensity range in the green channel for **(ii)**, **(iii)**, **(iv),** and **(v)** was rescaled with respect to that of **Figure 1E (i)** (lightest colour for pixels with 2 or more photons), for better visualization. Scale bar is 5 µm.

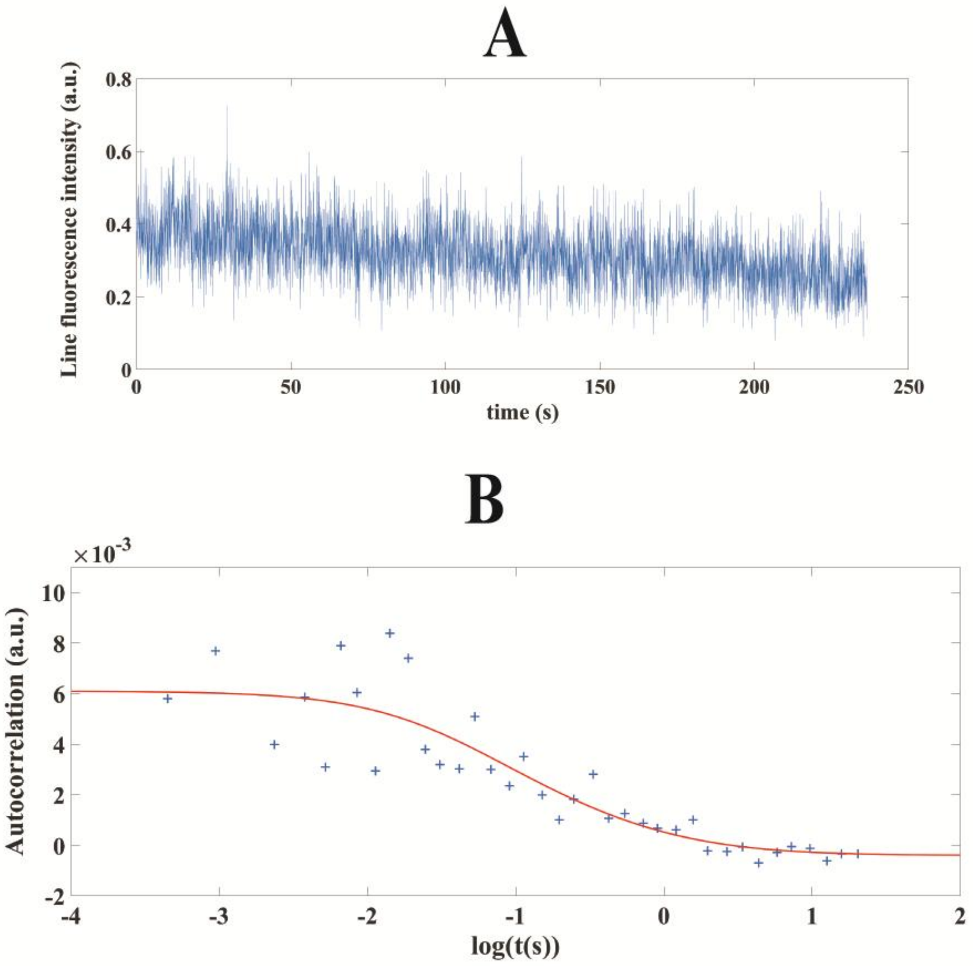

**Figure S8: Representative LsFCS data analysis for fluorescently labelled HCC bound on ALD.**
**A** shows a representative example of fluorescence intensity vs. time plots of lsFCS measurement of fluorescently labelled HCC on DOPC+PI(4)P ALDs at *pMMA* condition. **B** shows the corresponding autocorrelation curve, including a fit (solid line) to a 2D diffusion model.

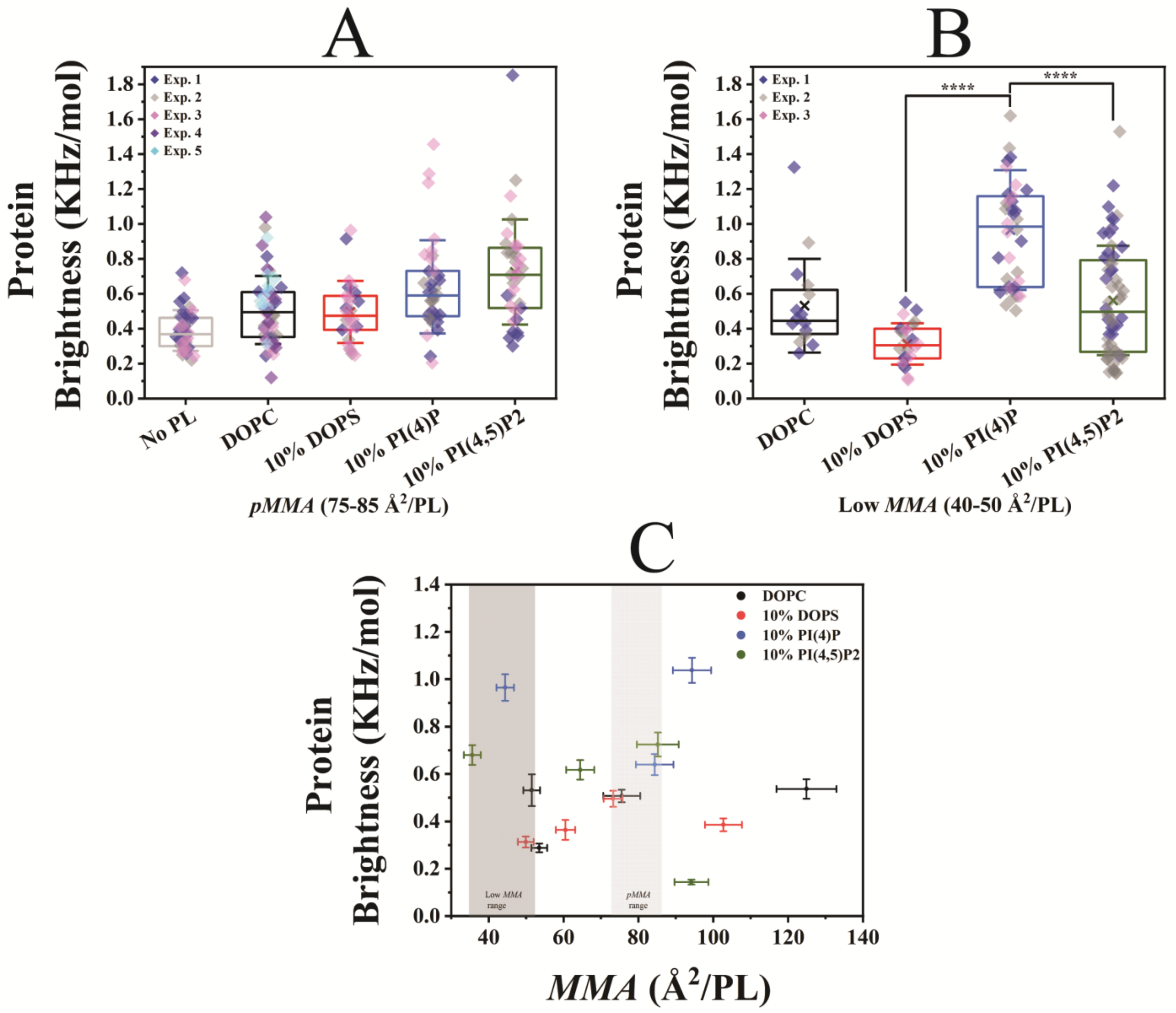


**Figure S9: HCC multimerization depends only weakly on ALD monolayer composition and *MMA*.**
Quantification of AF488-labelled HCC multimerization via lsFCS-based brightness analysis for the same ALD monolayers as in **Figures 1C**, **D**. **A** and **B** show the HCC brightness values for proteins binding to different ALD monolayers at *pMMA* and "low *MMA*", respectively, as box plots. The ALD monolayers are composed of DOPC, DOPC+DOPS (9:1), DOPC+PI(4)P (9:1) and DOPC+PI(4,5)P2 (9:1). **A** shows additionally the brightness data for ALDs made only of oil, i.e. bare oil droplets. Each point represents the result of one lsFCS measurement within one ALD. In all box plots, the horizontal line represents the median value, while the 'x' indicates the mean with the first and third quartile as the boundaries. The whiskers indicate SDs. Data points were pooled from at least two independent experiments denoted with individual colours. In **B**, statistical tests were performed via two-sample t-test at significance level 0.05 (**** $p<0.0001$). **C** shows the protein brightness values as a function of ALD monolayer *MMA*. *MMA* values were not directly measured but rather estimated from protein-free ALDs prepared in the same conditions, as indicated in **Table S6**. Light grey and dark grey boxes show the *pMMA* range and "low *MMA*" range, respectively. Here, each point represents the mean values of each of the two parameters obtained from at least 16-54 experimental points for DOPC ALDs, 26-36 points for 10% DOPS mixed ALDs, 35-38 points for 10% PI(4)P mixed ALDs and 32-64 points for 10% PI(4,5)P2 mixed ALDs with error bars as SEM. ALD monolayers were labelled with 0.001 molar% Rh-PE. See **Tables S3**, **S4** and **S6** for the details on the statistics.

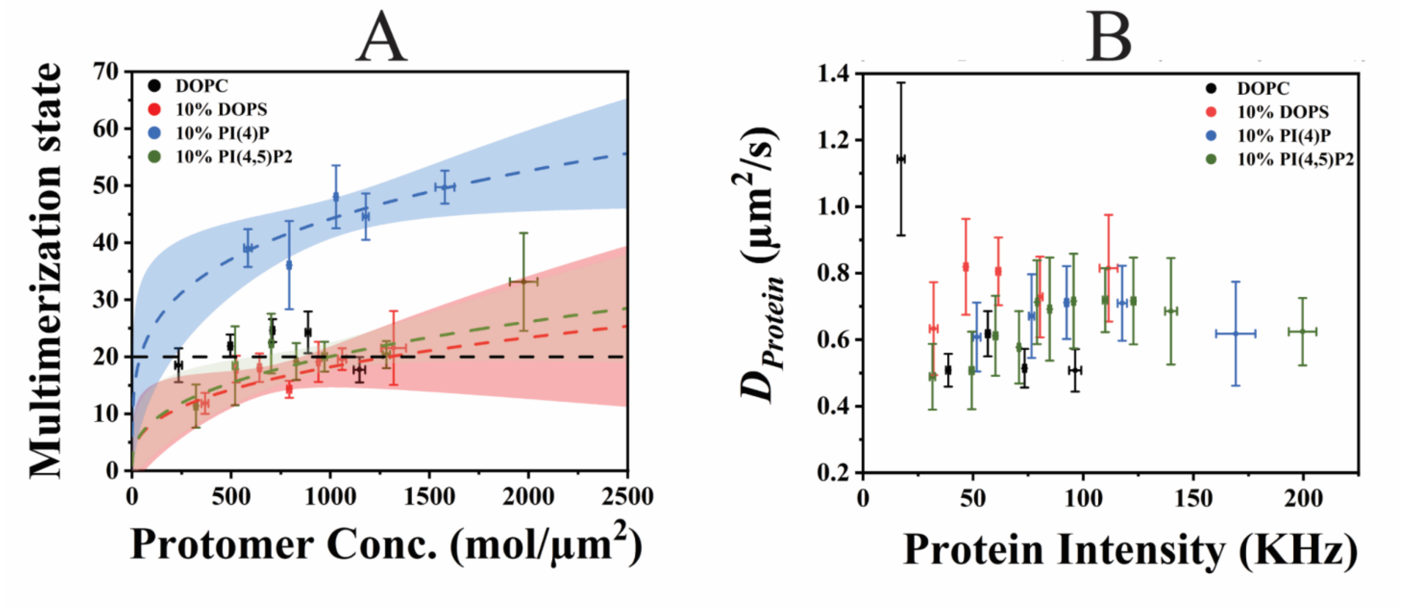


**Figure S10: HCC multimerization and mobility as a function of HCC concentration on ALDs.**
**A** shows the multimerization state values (via **Eq. 6**) plotted against the corresponding protein concentration (via **Eq. 5**). The data points correspond to those shown in **Figure 2**, with the same colour code. The dashed lines show the best fit to the experimental data points, using the empirical model *y=ax^b+1* (3). 95% confidence interval range for each fit is shown by the respective colour bands. The data for DOPC could not be fitted with the same model and, instead, only their average is shown as a dashed horizontal black line (i.e., *b* fixed to 0). The *a* value for 10% PI(4)P mixed ALDs is significantly higher than that for 10% DOPS and 10% PI(4,5)P2 mixed ALDs ($p<0.001$), as determined via one-way ANOVA at significance level 0.05. The *a* value difference between 10% DOPS and 10% PI(4,5)P2 mixed ALDs is not significant ($p>0.05$). **B** shows the protein diffusion coefficients $D_{Protein}$ for each LD composition as a function of the respective protein intensity values. As for **A**, data point binning was performed such that each point represents the mean value calculated from 18-37 points for DOPC LDs, 15-32 points for 10% DOPS mixed LDs, 13-35 points for 10% PI(4)P mixed LDs and 9-19 points for 10% PI(4,5)P2 mixed ALDs. The error bars in all figures represent the SEM calculated from the binning procedure. See **Table S6** for the details on the data points and fit parameters.

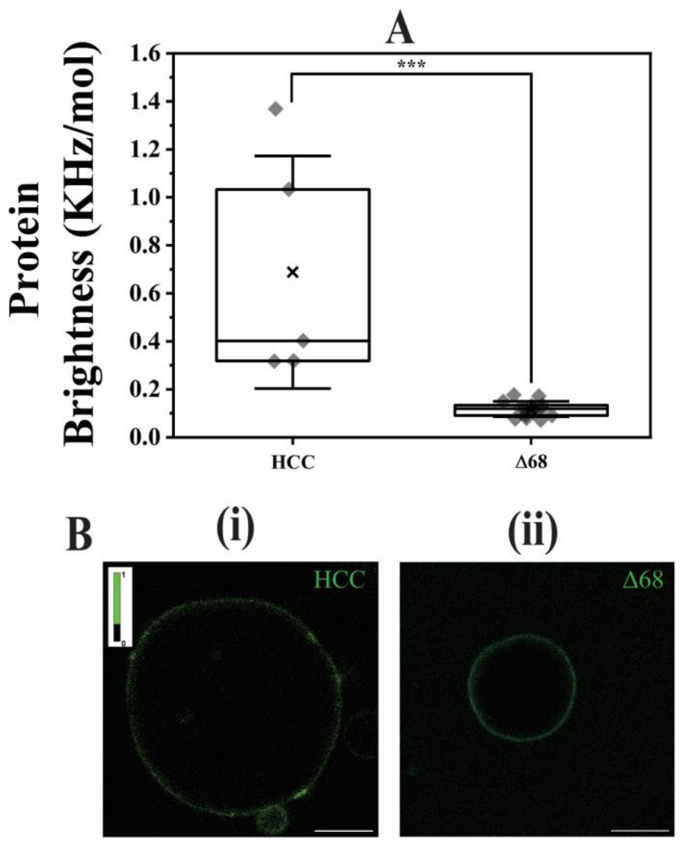

**Figure S11: Comparison of brightness between HCC and assembly-incompetent HCC mutant Δ68 on ALDs.** Quantification of protein (HCC and Δ68) multimerization was performed on the same type of ALD (*pMMA* condition) composed of DOPC+PI(4)P (9:1) using lsFCS to determine protein brightness (**A**). Protein concentration was similar (~10-20 KHz) for all measured samples. All values are shown as box plots. The horizontal lines mark the median, 'x' marks the mean with first and third quartile as the boundaries and whiskers as SD. Each data point represents a measurement from a single ALD. All statistical tests were performed via two-sample t-test at significance level 0.05 (*** $p<0.001$). See **Table S9** for the numerical values. **B** shows the fluorescence images of HCC- (or Δ68-) ALD interaction. The colour intensity range in the green channel for **(ii)**, **(iii)**, **(iv),** and **(v)** was rescaled with respect to that of **Figure 1E (i)** (lightest colour for pixels with 1 or more photons), for better visualization. Scale bar is 5 µm.

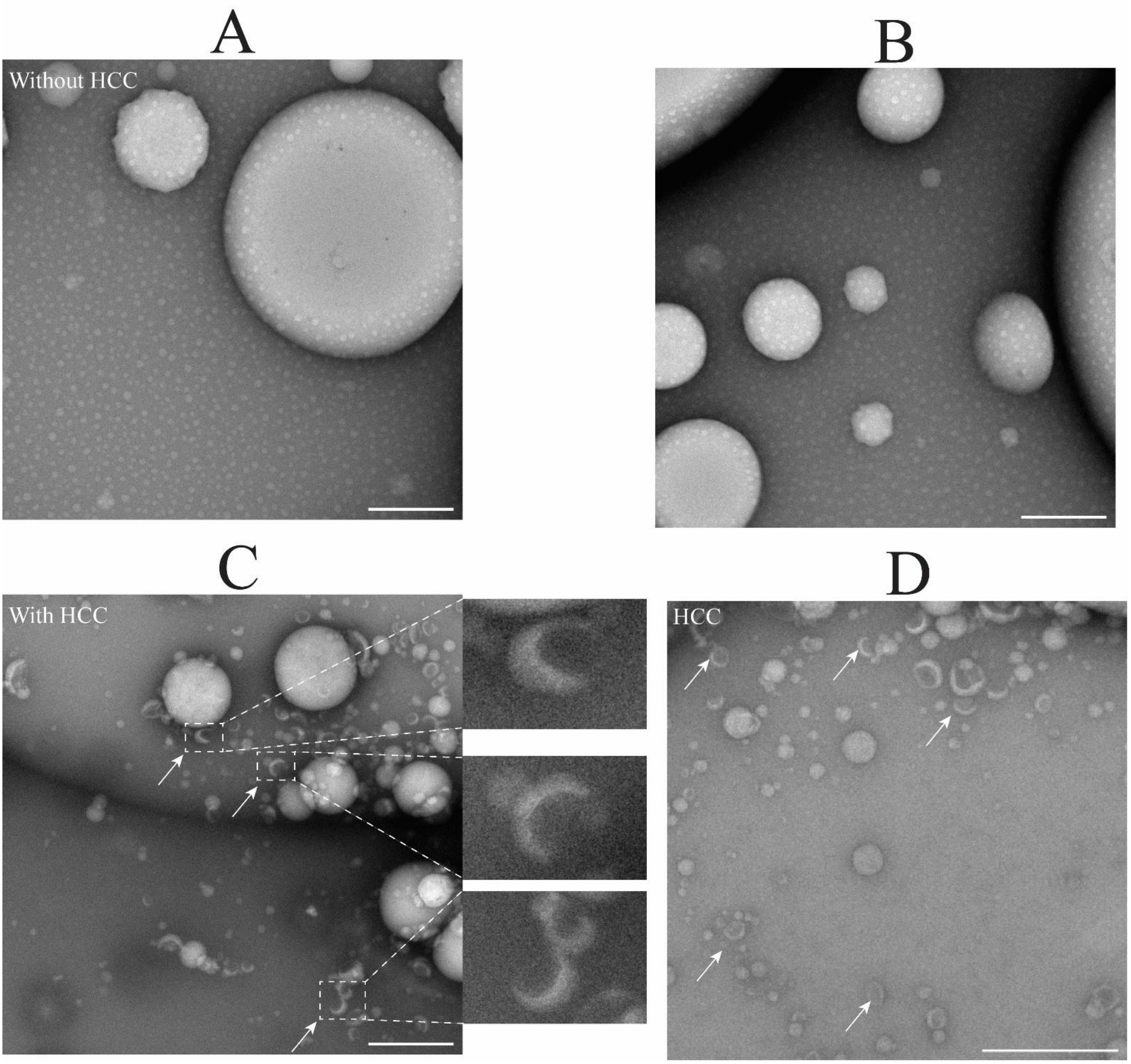


**Figure S12: TEM images of PI(4)P-containing ALDs with and without HCC.**
**A** and **B** show the TEM images of ALDs composed of DOPC+PI(4)P (9:1) monolayer, in the absence of protein. **C** shows the same in the presence of 200 nM HCC. Recurring protein structures in the form of crescents are indicated with white arrows. For better visualization, three such structures are also shown magnified on the right side of **C**. **D** shows a further example of the surface of an ALD, in the presence of HCC. Scale bar is 500 nm.

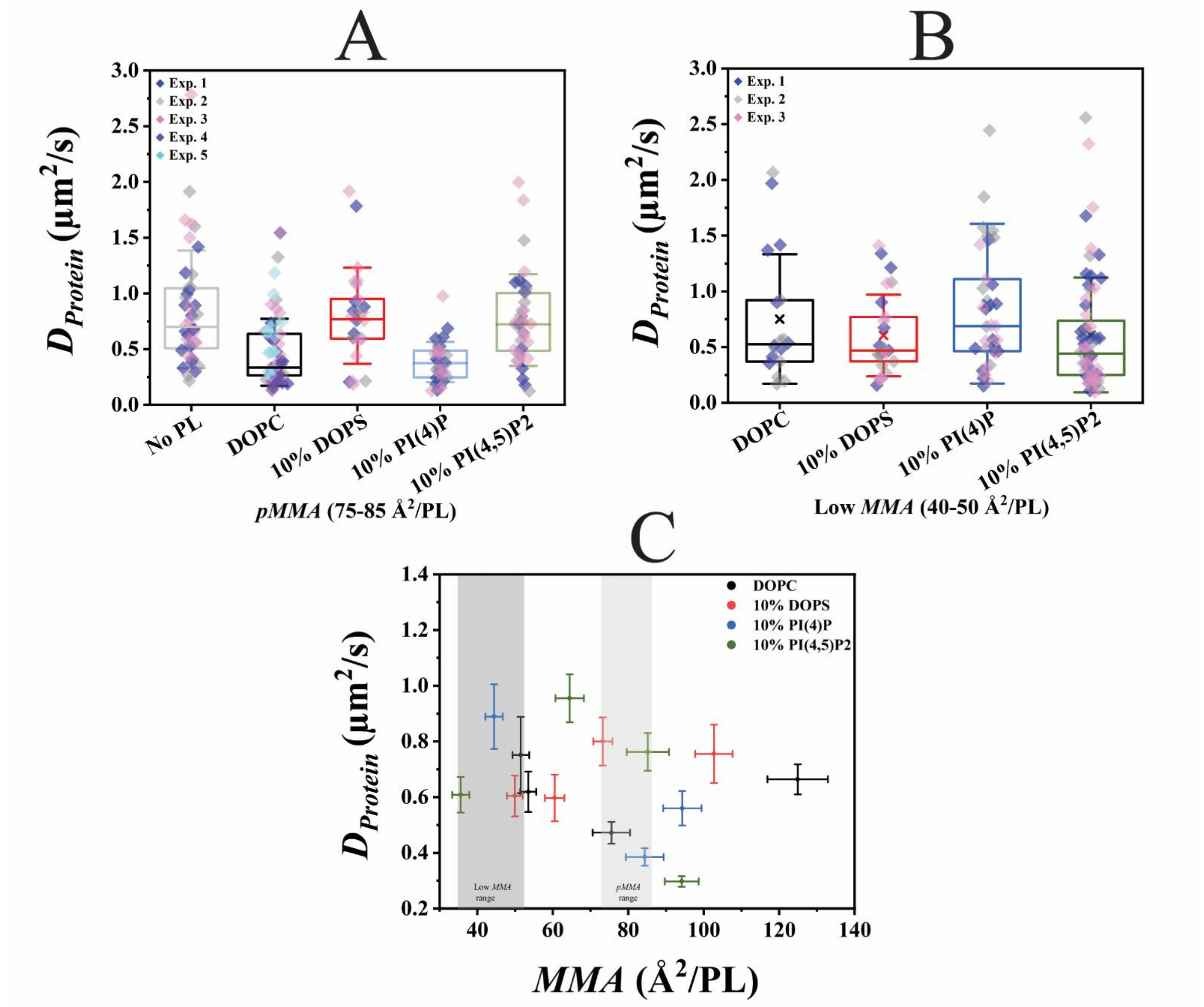


**Figure S13: HCC mobility does not strongly depend on ALD monolayer composition and *MMA*.**
AF488-labelled HCC mobility was quantified via lsFCS for the same ALD monolayers as in **Figures 1C** and **D** and represented here as protein diffusion coefficient ($D_{Protein}$). **A** and **B** represent the distribution of $D_{Protein}$ values for proteins bound to ALD monolayers at *pMMA* and "low *MMA"* conditions, respectively. The ALD monolayers were composed of DOPC, DOPC+DOPS (9:1), DOPC+PI(4)P (9:1) and DOPC+PI(4,5)P2 (9:1). **A** additionally shows $D_{Protein}$ data for ALDs made only with oil, i.e. bare oil droplets. Each point represents the result of one lsFCS measurement within one ALD. In all box plots, the horizontal line represents the median value, while the 'x' indicates the mean with the first and third quartile as the boundaries. The whiskers indicate SDs. Data points were pooled from at least two independent experiments denoted with individual colours. **C** shows the $D_{Protein}$ values as a function of ALD monolayer *MMA*. *MMA* values were not directly measured but rather estimated from PL/oil ratios and protein-free ALDs, as indicated on **Table S6**. Light grey and dark grey boxes show the *pMMA* range and "low *MMA*" range respectively. Here, each point represents the mean values of each of the two parameters obtained from at least 18-59 experimental points for DOPC ALDs, 25-37 points for 10% DOPS mixed ALDs, 33-38 points for 10% PI(4)P mixed ALDs and 32-64 points for 10% PI(4,5)P2 mixed ALDs with error bars as SEM. ALD monolayers were labelled with 0.001 molar% Rh-PE. See **Tables S3**, **S4** and **S6** for the details on the statistics.

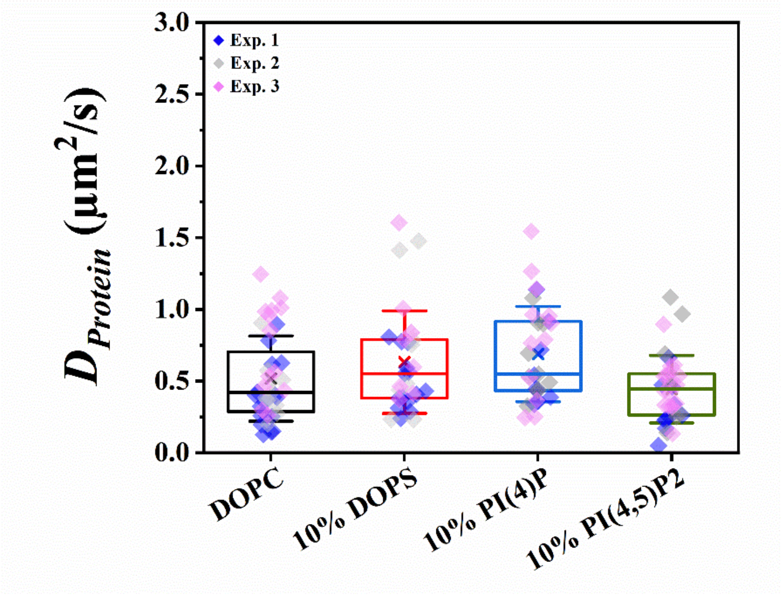


**Figure S14: HCC mobility on ALD monolayers at *pMMA* is not strongly affected by reducing environment.** Quantification of AF488 labelled HCC mobility via lsFCS in reducing environment on the same ALD monolayers as in **Figure 3**, represented as protein diffusion coefficient ($D_{Protein}$). $D_{Protein}$ values for HCC associated to ALD monolayers composed of DOPC, DOPC+DOPS (9:1), DOPC+PI(4)P (9:1) and DOPC+PI(4,5)P2 (9:1) presented as box plots. In all box plots, the horizontal line is the median, 'x' marks the mean with first and third quartile as the boundaries and whiskers as SD. All data points were obtained from three independent experiments denoted with individual colours. Reducing environment was obtained through the addition of 5 % β-mercaptoethanol (v/v). See Table **S5** for the numerical values.

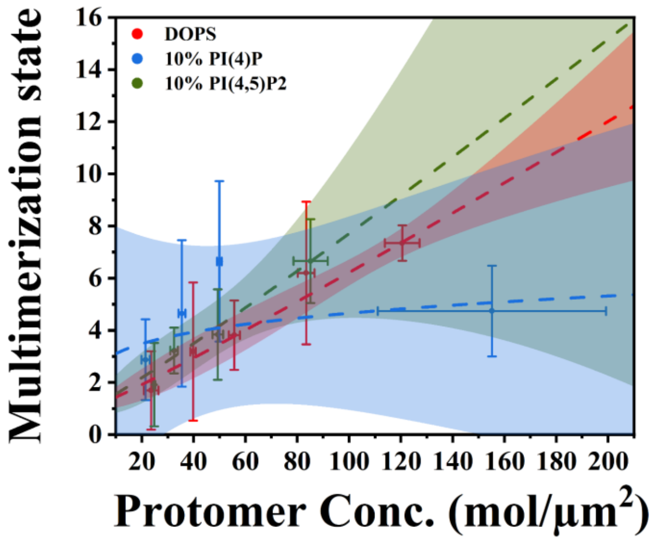


**Figure S15: HCC limited oligomerization as a function of its concentration on GUVs.** Protein concentration and the corresponding multimerization state data points from **Figure 4** were plotted relative to each other. Protein (protomer) concentration was obtained from protein intensity values and multimerization state was obtained from the corresponding brightness values using Eqs. 5 and 6, respectively. As an approximation, brightness value for the lowest multimerization state (possibly, monomer) $B_{monomer}$ was assumed to be 0.06. Fitting was performed is the same way as for **Figure S10A**. 95% confidence interval range for each fit is shown by the respective colour bands. For a better visualization, the data points were binned such that each point in the graph represents the mean value calculated from 5-12 points for 10% DOPS mixed GUVs, 5-10 points for 10% PI(4)P mixed LDs and 5-11 points for 10% PI(4,5)P2 mixed LDs. Error bars are SEM calculated from the binning procedure. The *a* value difference between 10% DOPS mixed ALDs, 10% PI(4)P mixed ALDs and 10% PI(4,5)P2 mixed ALDs is not significant ($p>0.05$) as determined via one-way ANOVA at significance level 0.05. See **Table S8** for the data points and fit parameters.

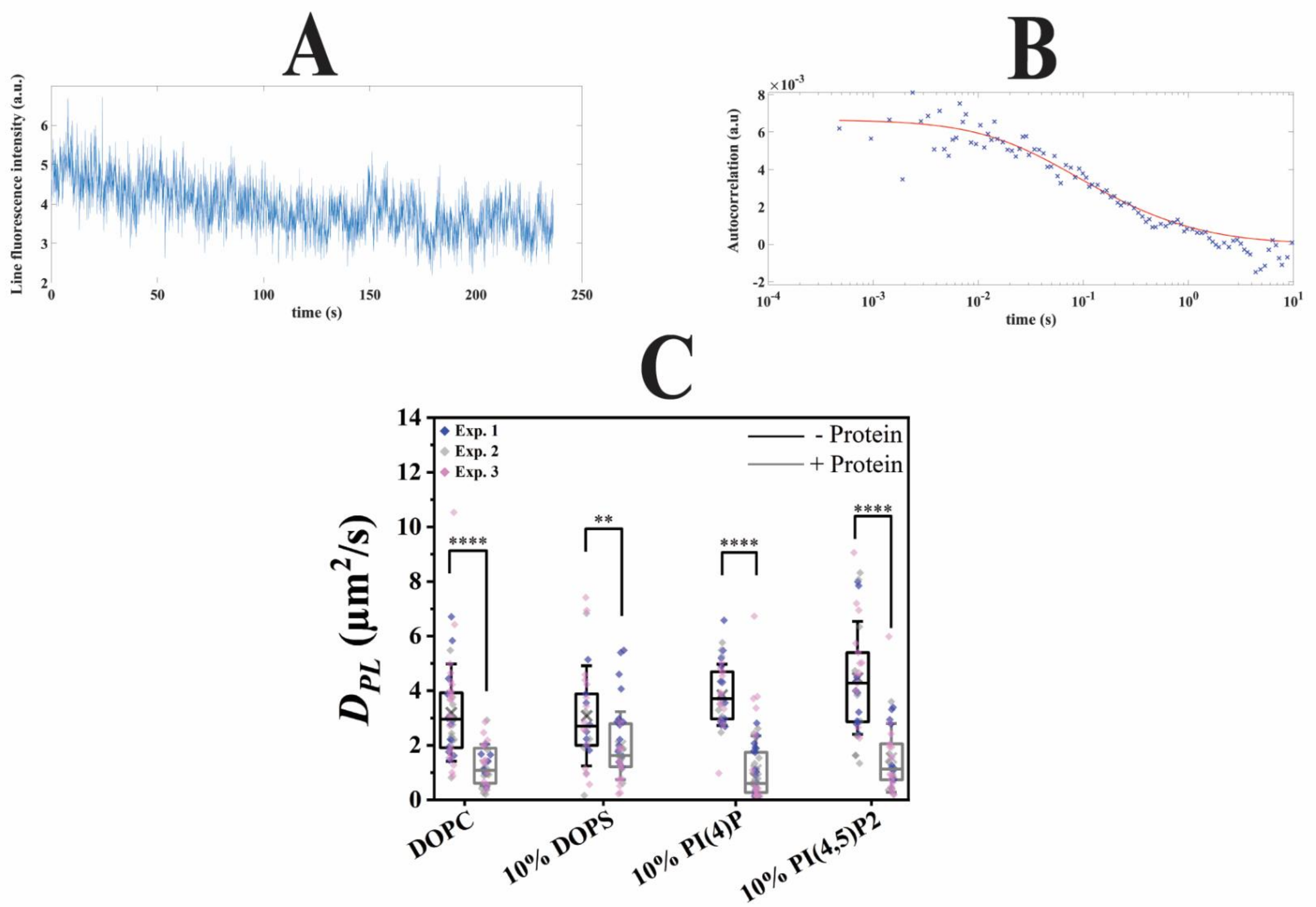


**Figure S16: Effect of HCC binding on monolayer PL mobility measured using TF-PC.**
**A** shows a representative example of fluorescence intensity vs. time plots of lsFCS measurement of fluorescently labelled lipids (Rh-PE) in the presence of HCC, for DOPC+PI(4)P ALDs at *pMMA* condition. **B** shows the corresponding autocorrelation curve, including a fit (solid line) to a 2D diffusion model. C shows the PL mobility on ALDs composed of DOPC, DOPC+DOPS (9:1), DOPC+PI(4)P (9:1) and DOPC+PI(4,5)P2 (9:1) at *pMMA* (labelled with 0.005 molar% TF-PC), represented as $D_{PL/TF\text{-}PC}$ was analysed by lsFCS. All values are shown as box plots with the data points obtained from three independent experiments denoted with individual colours. The data points for ALDs without HCC are depicted in black boxes and those in the presence of HCC in grey boxes. In all box plots, the horizontal line marks the median, ‘x’ marks the mean with first and third quartile as the boundaries and whiskers as SD. All statistical tests were performed via two-sample t-test at significance level 0.05 (**** $p<0.0001$ and ** $p<0.01$). See **Table S10** for the numerical values.

**Supplemental tables**

**Table S1: Normalized binding intensity of HCC on PIP™ strip related to Figures 1B and S1**

| **PL spot** | **Normalized protein intensity** (a.u) (n=3) Mean ± SD |
|---|---|
| PI(4)P | 1.00 ± 0.09 |
| PI(3)P | 0.95 ± 0.20 |
| PI(5)P | 0.89 ± 0.22 |
| PI(3,4,5)P3 | 0.9 ± 0.3 |
| PI(3,4)P2 | 0.4 ± 0.4 |
| PI(3,5)P2 | 0.71 ± 0.24 |
| PI(4,5)P2 | 0.8 ± 0.3 |
| PA | 0.71 ± 0.28 |
| PI | 0.6 ± 0.5 |
| PS | 0.76 ± 0.10 |
| PE | 0.4 ± 0.3 |
| PC | 0.20 ± 0.09 |

**Table S2: Mean values of initial fluorescently labelled protein intensity during the course of a typical measurement related to Figure S4**

| **ALD number** | **Protein intensity** (KHz) (n=3) Mean ± SEM |
|---|---|
| ALD 1 | 58.6 ± 10.6 |
| ALD 2 | 45.8 ± 11.4 |
| ALD 3 | 49.0 ± 14.3 |
| ALD 4 | 64.30 ± 3.88 |
| ALD 5 | 69.0 ± 18.3 |
| ALD 6 | 50.2 ± 6.07 |
| ALD 7 | 71.8 ± 27.7 |
| ALD 8 | 47.1 ± 17.3 |

**Table S3: HCC parameters measured by lsFCS on ALD monolayers under *pMMA* conditions (75-85 Å$^2$/PL) related to Figures 1C, S6A, S9A and S13A**

| **Parameter** | **No PL** | **DOPC LD** (*pMMA*) | **DOPC+DOPS LD** (9:1 molar ratio) (*pMMA*) | **DOPC+PI(4)P LD** (9:1 molar ratio) (*pMMA*) | **DOPC+PI(4,5)P2 LD** (9:1 molar ratio) (*pMMA*) | **DOPC+PI(4)P LD** (9.66:0.34 molar ratio) (*pMMA*) | **DOPC+PI(4,5)P2 LD** (9.77:0.23 molar ratio) (*pMMA*) |
|---|---|---|---|---|---|---|---|
| Protein Intensity (KHz) Mean ± SD | 124 ± 31 (n=60) | 62 ± 22 (n=59) | 71 ± 25 (n=44) | 90 ± 30 (n=37) | 90 ± 40 (n=44) | 80 ± 30 (n=32) | 100 ± 50 (n=22) |
| Protein Brightness (KHz/mol) Mean ± SD | 0.400 ± 0.117 (n=40) | 0.50 ± 0.20 (n=54) | 0.50 ± 0.20 (n=27) | 0.64 ± 0.30 (n=36) | 0.72 ± 0.30 (n=35) | | |
| $D_{Protein}$ (μm$^2$/s) Mean ± SD | 0.9 ± 0.5 (n=40) | 0.5 ± 0.3 (n=59) | 0.8 ± 0.4 (n=25) | 0.4 ± 0.2 (n=33) | 0.8 ± 0.4 (n=37) | | |

**Table S4: HCC parameters measured by lsFCS on ALD monolayers under highly compressed conditions (40-50 Å$^2$/PL) related to Figures 1D, S6B, S9B and S13B**

| **Parameter** | **DOPC LD** | **DOPC+DOPS LD** (9:1 molar ratio) | **DOPC+PI(4)P LD** (9:1 molar ratio) | **DOPC+PI(4,5)P2 LD** (9:1 molar ratio) | **DOPC+PI(4)P LD** (9.66:0.34 molar ratio) | **DOPC+PI(4,5)P2 LD** (9.77:0.23 molar ratio) |
|---|---|---|---|---|---|---|
| Protein Intensity (KHz) Mean ± SD | 24 ± 15 (n=18) | 55 ± 23 (n=28) | 96 ± 26 (n=38) | 120 ± 50 (n=69) | 90 ± 40 (n=37) | 80 ± 40 (n=30) |
| Protein Brightness (KHz/mol) Mean ± SD | 0.5 ± 0.3 (n=16) | 0.30 ± 0.10 (n=26) | 1 ± 0.3 (n=38) | 0.6 ± 0.3 (n=64) | | |
| $D_{Protein}$ (μm$^2$/s) Mean ± SD | 0.8 ± 0.6 (n=18) | 0.8 ± 0.4 (n=25) | 0.9 ± 0.7 (n=38) | 0.6 ± 0.5 (n=64) | | |

**Table S5: HCC parameters (reduced environment) measured by lsFCS on ALD monolayers under *pMMA* conditions (75-85 Å$^2$/PL) related to Figures 3 and S14**

**i)**

| **Parameter** | **DOPC LD** (*pMMA*) | **DOPC+DOPS LD** (9:1 molar ratio) (*pMMA*) | **DOPC+PI(4)P LD** (9:1 molar ratio) (*pMMA*) | **DOPC+PI(4,5)P2 LD** (9:1 molar ratio) (*pMMA*) |
|---|---|---|---|---|
| Protein Intensity (KHz) Mean ± SD | 7.3 ± 2.4 (n=45) | 9.4 ± 2.6 (n=34) | 19.00 ± 11.22 (n=36) | 14 ± 7 (n=34) |
| Protein Brightness (KHz/mol) Mean ± SD | 0.12 ± 0.06 (n=42) | 0.50 ± 0.22 (n=33) | 0.7 ± 0.3 (n=35) | 0.40 ± 0.21 (n=34) |
| $D_{Protein}$ (µm$^2$/s) Mean ± SD | 0.6 ± 0.6 (n=42) | 0.6 ± 0.4 (n=31) | 0.7 ± 0.3 (n=31) | 0.44 ± 0.23 (n=34) |

**ii) Data points related to Figure 3D, DOPC**

| **Parameter** | **Point 1** | **Point 2** | **Point 3** | **Point 4** | **Point 5** | **Point 6** |
|---|---|---|---|---|---|---|
| Protein Intensity (KHz) Mean ± SEM | 4.12 ± 0.487 (n=10) | 5.98 ± 0.0820 (n=5) | 6.88 ± 0.123 (n=5) | 7.94 ± 0.154 (n=10) | 9.16 ± 0.137 (n=7) | 11.20 ± 0.647 (n=5) |
| Protein Concentration (mol/µm$^2$) Mean ± SEM | 44 ± 5 (n=10) | 64 ± 1 (n=5) | 73 ± 1 (n=5) | 85 ± 2 (n=10) | 98 ± 2 (n=7) | 120 ± 7 (n=5) |
| Protein Brightness (KHz/mol) Mean ± SEM | 0.151 ± 0.0254 (n=10) | 0.145 ± 0.0112 (n=5) | 0.103 ± 0.0157 (n=5) | 0.0968 ± 0.0119 (n=10) | 0.134 ± 0.0388 (n=7) | 0.113 ± 0.0217 (n=5) |
| Multimerization state Mean ± SEM | 3.75 ± 1.31 (n=10) | 3.44 ± 0.576 (n=5) | 1.30 ± 0.808 (n=5) | 0.983 ± 0.616 (n=10) | 2.92 ± 1.990 (n=7) | 1.81 ± 1.12 (n=5) |

**ii) Data points related to Figure 3D, DOPC+10%DOPS (9:1)**

| **Parameter** | **Point 1** | **Point 2** | **Point 3** | **Point 4** | **Point 5** |
|---|---|---|---|---|---|
| Protein Intensity (KHz) Mean ± SEM | 5.93 ± 0.373 (n=5) | 8.13 ± 0.119 (n=5) | 8.70 ± 0.0566 (n=8) | 9.85 ± 0.331 (n=9) | 13.8 ± 0.821 (n=6) |
| Protein Concentration (mol/µm$^2$) Mean ± SEM | 63 ± 4 (n=5) | 87 ± 1 (n=5) | 93 ± 1 (n=8) | 105 ± 4 (n=9) | 147 ± 9 (n=6) |
| Protein Brightness (KHz/mol) | 0.433 ± 0.0897 | 0.489 ± 0.129 | 0.459 ± 0.0616 | 0.550 ± 0.101 | 0.555 ± 0.0560 |

| | | | | | |
|---|---|---|---|---|---|
| Mean ± SEM | (n=5) | (n=5) | (n=8) | (n=9) | (n=6) |
| Multimerization state Mean ± SEM | 18.3 ± 4.62 (n=5) | 21.2 ± 6.69 (n=5) | 19.7 ± 3.17 (n=8) | 24.3 ± 5.22 (n=9) | 24.6 ± 2.89 (n=6) |

**iii) Data points related to Figure 3D, DOPC+10%PI(4)P (9:1)**

| **Parameter** | **Point 1** | **Point 2** | **Point 3** | **Point 4** | **Point 5** | **Point 6** |
|---|---|---|---|---|---|---|
| Protein Intensity (KHz) Mean ± SEM | 9.98 ± 0.499 (n=10) | 14.4 ± 0.219 (n=6) | 16.7 ± 0.4416 (n=6) | 20.6 ± 0.499 (n=7) | 25.3 ± 0.541 (n=3) | 43.9 ± 8.08 (n=4) |
| Protein Concentration (mol/μm²) Mean ± SEM | 106 ± 5 (n=10) | 153 ± 2 (n=6) | 178 ± 5 (n=6) | 219 ± 5 (n=7) | 269 ± 6 (n=3) | 467 ± 90 (n=4) |
| Protein Brightness (KHz/mol) Mean ± SEM | 0.678 ± 0.121 (n=10) | 0.764 ± 0.149 (n=6) | 0.867 ± 0.157 (n=6) | 0.510 ± 0.0424 (n=7) | 0.793 ± 0.284 (n=3) | 0.937 ± 0.148 (n=4) |
| Multimerization state Mean ± SEM | 31 ± 6 (n=10) | 35 ± 8 (n=6) | 41 ± 8 (n=6) | 22 ± 2 (n=7) | 37 ± 15 (n=3) | 44 ± 8 (n=4) |

**iv) Data points related to Figure 3D, DOPC+10%PI(4,5)P2 (9:1)**

| **Parameter** | **Point 1** | **Point 2** | **Point 3** | **Point 4** | **Point 5** | **Point 6** |
|---|---|---|---|---|---|---|
| Protein Intensity (KHz) Mean ± SEM | 6.38 ± 1.12 (n=4) | 10.0 ± 0.265 (n=6) | 11.0 ± 0.219 (n=3) | 12.3 ± 0.248 (n=5) | 13.9 ± 0.217 (n=6) | 20.7 ± 2.58 (n=10) |
| Protein Concentration (mol/μm²) Mean ± SEM | 68 ± 10 (n=4) | 106 ± 3 (n=6) | 117 ± 2 (n=3) | 130 ± 3 (n=5) | 148 ± 2 (n=6) | 220 ± 27 (n=10) |
| Protein Brightness (KHz/mol) Mean ± SEM | 0.310 ± 0.0419 (n=4) | 0.465 ± 0.136 (n=6) | 0.365 ± 0.0612 (n=3) | 0.377 ± 0.0696 (n=5) | 0.404 ± 0.0665 (n=6) | 0.429 ± 0.0788 (n=10) |
| Multimerization state Mean ± SEM | 11.9 ± 2.16 (n=4) | 20 ± 7 (n=6) | 14.8 ± 3.15 (n=3) | 15 ± 4 (n=5) | 16.8 ± 3.42 (n=6) | 18 ± 4 (n=10) |

**v) Fit parameters connected to Figure 3D**

| **Parameter** | **DOPC** | **DOPC+DOPS** (9:1 molar ratio) | **DOPC+PI(4)P** (9:1 molar ratio) | **DOPC+PI(4,5)P2** (9:1 molar ratio) |
|---|---|---|---|---|
| *a* Mean ± SEM | 1 ± 10 (n=6) | 3.4 ± 1.7 (n=5) | 11.2 ± 3.6 (n=6) | 2.25 ± 1.08 (n=6) |
| *b* Mean ± SEM | 0 ± 2 (n=6) | 0.389 ± 0.106 (n=5) | 0.223 ± 0.0602 (n=6) | 0.384 ± 0.0990 (n=6) |

**Table S6: Parameters as a function of *MMA* related to Figures S2, S3, S5, S9C and S13C**

**i) DOPC ALD**

| PL: Oil mass ratio | 1:2000 | 1:1000 | 1:500 | 1:250 |
|---|---|---|---|---|
| *MMA*<br>(Å$^2$/PL)<br>Mean ± SEM | 125 ± 8<br>(n=47) | 76 ± 5<br>(n=29) | 53.5 ± 2.1<br>(n=42) | 51.5 ± 2.2<br>(n=51) |
| $D_{PL}$<br>(μm$^2$/s)<br>Mean ± SEM | 8.96 ± 0.20<br>(n=47) | 7.66 ± 0.10<br>(n=29) | 6.13 ± 0.12<br>(n=42) | 5.84 ± 0.12<br>(n=51) |
| TF-PC intensity<br>(a.u) ($10^4$)<br>Mean ± SD | | 2.2 ± 0.4<br>(n=9) | | |
| TF-PI(4)P intensity<br>(a.u) ($10^4$)<br>Mean ± SD | | 1.9 ± 0.6<br>(n=13) | | |
| TF-PI(4,5)P 2<br>intensity (a.u) ($10^4$)<br>Mean ± SD | | 2.2 ± 0.4<br>(n=16) | | |
| Protein Intensity<br>(KHz)<br>Mean ± SEM | 63.4 ± 2.6<br>(n=49) | 61.7 ± 2.9<br>(n=59) | 41 ± 4<br>(n=35) | 24 ± 3<br>(n=18) |
| Protein Brightness<br>(KHz/mol)<br>Mean ± SEM | 0.54 ± 0.04<br>(n=37) | 0.507 ± 0.027<br>(n=54) | 0.288 ± 0.018<br>(n=30) | 0.53 ± 0.07<br>(n=16) |
| $D_{Protein}$ (μm$^2$/s)<br>Mean ± SEM | 0.66 ± 0.05<br>(n=41) | 0.47 ± 0.04<br>(n=59) | 0.62 ± 0.07<br>(n=30) | 0.75 ± 0.14<br>(n=18) |

**ii) DOPC + DOPS (9:1 molar ratio) ALD**

| PL: Oil mass ratio | 1:2000 | 1:1000 | 1:500 | 1:250 |
|---|---|---|---|---|
| *MMA*<br>(Å$^2$/PL)<br>Mean ± SEM | 103 ± 5<br>(n=21) | 73.2 ± 2.5<br>(n=42) | 60.5 ± 2.6<br>(n=38) | 49.9 ± 2.1<br>(n=38) |
| $D_{PL}$<br>(μm$^2$/s)<br>Mean±SEM | 8.3 ± 0.4<br>(n=21) | 6.53 ± 0.23<br>(n=42) | 4.91 ± 0.14<br>(n=38) | 5.01 ± 0.15<br>(n=38) |
| Protein Intensity<br>(KHz)<br>Mean ± SEM | 76.7 ± 3.0<br>(n=51) | 71 ± 4<br>(n=44) | 57 ± 4<br>(n=34) | 55 ± 4<br>(n=28) |
| Protein Brightness<br>(KHz/mol)<br>Mean ± SEM | 0.385 ± 0.027<br>(n=36) | 0.50 ± 0.03<br>(n=27) | 0.40 ± 0.04<br>(n=26) | 0.313 ± 0.023<br>(n=26) |
| $D_{Protein}$ (μm$^2$/s)<br>Mean ± SEM | 0.76 ± 0.10<br>(n=37) | 0.80 ± 0.09<br>(n=25) | 0.60 ± 0.08<br>(n=25) | 0.60 ± 0.07<br>(n=25) |

**iii) DOPC + PI(4)P (9:1 molar ratio) ALD**

| PL: Oil mass ratio | 1:1000 | 1:500 | 1:250 |
|---|---|---|---|
| *MMA*<br>(Å$^2$/PL)<br>Mean ± SEM | 94 ± 5<br>(n=46) | 84 ± 5<br>(n=44) | 44.24 ± 2.37<br>(n=47) |
| $D_{PL}$<br>(μm$^2$/s)<br>Mean±SEM | 6.01 ± 0.17<br>(n=46) | 5.60 ± 0.21<br>(n=44) | 4.94 ± 0.16<br>(n=47) |
| Protein Intensity<br>(KHz)<br>Mean ± SEM | 95.90 ± 7.50<br>(n=40) | 91.32 ± 5.52<br>(n=37) | 95.81 ± 4.20<br>(n=38) |

| | | | |
|---|---|---|---|
| Protein Brightness (KHz/mol) Mean ± SEM | 1.04 ± 0.0500 (n=35) | 0.640 ± 0.0400 (n=36) | 0.970 ± 0.0600 (n=38) |
| $D_{Protein}$ (μm²/s) Mean ± SEM | 0.560 ± 0.0600 (n=33) | 0.390 ± 0.0300 (n=33) | 0.890 ± 0.120 (n=38) |

**iv) DOPC + PI(4,5)P2 (9:1 molar ratio) ALD**

| PL: Oil mass ratio | 1:2000 | 1:1000 | 1:500 | 1:250 |
|---|---|---|---|---|
| *MMA* (Å²/PL) Mean ± SEM | 94.14 ± 4.50 (n=33) | 85.2 ± 5.6 (n=45) | 64.4 ± 3.8 (n=24) | 35.60 ± 2.27 (n=53) |
| $D_{PL}$ (μm²/s) Mean±SEM | 5.60 ± 0.18 (n=33) | 5.34 ± 0.18 (n=45) | 5.18 ± 0.25 (n=24) | 4.83 ± 0.14 (n=53) |
| Protein Intensity (KHz) Mean ± SEM | 65 ± 4 (n=57) | 93 ± 7 (n=44) | 94 ± 6 (n=38) | 116 ± 6 (n=69) |
| Protein Brightness (KHz/mol) Mean ± SEM | 0.14 ± 0.01 (n=55) | 0.72 ± 0.05 (n=35) | 0.62 ± 0.04 (n=32) | 0.60 ± 0.04 (n=64) |
| $D_{Protein}$ (μm²/s) Mean ± SEM | 0.297 ± 0.019 (n=52) | 0.76 ± 0.07 (n=37) | 0.95 ± 0.09 (n=32) | 0.61 ± 0.06 (n=64) |

**Table S7:**

**(i) DOPC ALD data points related to Figures 2 and S10A**

| Parameter | Point 1 | Point 2 | Point 3 | Point 4 | Point 5 |
|---|---|---|---|---|---|
| Protein Intensity (KHz) Mean ± SEM | 22.0 ± 1.6 (n=37) | 46.5 ± 0.9 (n=43) | 66.5 ± 0.9 (n=38) | 83.37 ± 1.19 (n=18) | 107.7 ± 2.7 (n=9) |
| Protein Concentration (mol/μm²) Mean ± SEM | 234.32 ± 17.1 (n=37) | 495 ± 10 (n=43) | 708 ± 10 (n=38) | 887.3 ± 12.7 (n=18) | 1146.50 ± 29.13 (n=9) |
| Protein Brightness (KHz/mol) Mean ± SEM | 0.44 ± 0.06 (n=37) | 0.50 ± 0.04 (n=43) | 0.56 ± 0.04 (n=38) | 0.55 ± 0.07 (n=18) | 0.42 ± 0.04 (n=9) |
| Multimerization state Mean ± SEM | 19 ± 3 (n=37) | 22 ± 2 (n=43) | 25 ± 2 (n=38) | 24 ± 4 (n=18) | 18.0 ± 2.2 (n=9) |

**(ii) DOPC+DOPS (9:1) ALD data points related to Figures 2 and S10A**

| Parameter | Point 1 | Point 2 | Point 3 | Point 4 | Point 5 | Point 6 | Point 7 |
|---|---|---|---|---|---|---|---|
| Protein Intensity (KHz) Mean ± SEM | 34.5 ± 1.7 (n=20) | 49.3 ± 0.7 (n=20) | 60.4 ± 0.8 (n=20) | 74 ± 1 (n=28) | 88.3 ± 0.6 (n=10) | 100 ± 2 (n=11) | 124 ± 6 (n=10) |
| Protein Concentration (mol/μm²) Mean ± SEM | 367 ± 18 (n=20) | 525 ± 8 (n=20) | 643 ± 8 (n=20) | 792 ± 10 (n=28) | 940 ± 7 (n=10) | 1059 ± 21 (n=11) | 1318 ± 63 (n=10) |

| | | | | | | | |
|---|---|---|---|---|---|---|---|
| Protein Brightness (KHz/mol) Mean ± SEM | 0.31 ± 0.04 (n=20) | 0.42 ± 0.05 (n=20) | 0.43 ± 0.05 (n=20) | 0.36 ± 0.03 (n=28) | 0.45 ± 0.07 (n=10) | 0.46 ± 0.04 (n=11) | 0.496 ± 0.126 (n=10) |
| Multimerization state Mean ± SEM | 11.8 ± 1.8 (n=20) | 17.8 ± 2.3 (n=20) | 18.1 ± 2.5 (n=20) | 14.3 ± 1.5 (n=28) | 19 ± 4 (n=10) | 19.6 ± 1.9 (n=11) | 21.5 ± 6.5 (n=10) |

**(iii) DOPC+PI(4)P (9:1) ALD data points related to Figures 2 and S10A**

| Parameter | Point 1 | Point 2 | Point 3 | Point 4 | Point 5 |
|---|---|---|---|---|---|
| Protein Intensity (KHz) Mean ± SEM | 54.9 ± 1.8 (n=30) | 74.4 ± 0.8 (n=7) | 96.6 ± 0.7 (n=13) | 110.7 ± 1.5 (n=24) | 148.2 ± 4.5 (n=23) |
| Protein Concentration (mol/µm$^2$) Mean ± SEM | 584 ± 19 (n=30) | 792 ± 8 (n=7) | 1028 ± 8 (n=13) | 1179 ± 16 (n=24) | 1580 ± 50 (n=23) |
| Protein Brightness (KHz/mol) Mean ± SEM | 0.84 ± 0.06 (n=30) | 0.78 ± 0.15 (n=7) | 1.01 ± 0.11 (n=13) | 0.94 ± 0.08 (n=24) | 1.04 ± 0.06 (n=23) |
| Multimerization state Mean ± SEM | 39 ± 3 (n=30) | 36 ± 8 (n=7) | 48 ± 6 (n=13) | 45 ± 4 (n=24) | 50 ± 3 (n=23) |

**(iv) DOPC+PI(4,5)P2 (9:1) ALD data points related to Figures 2 and S10A**

| Parameter | Point 1 | Point 2 | Point 3 | Point 4 | Point 5 | Point 6 | Point 7 |
|---|---|---|---|---|---|---|---|
| Protein Intensity (KHz) Mean ± SEM | 29.2 ± 1.0 (n=13) | 48.8 ± 1.2 (n=31) | 66.4 ± 0.8 (n=26) | 77.8 ± 0.5 (n=22) | 91.4 ± 1.1 (n=36) | 120.5 ± 1.6 (n=37) | 186 ± 7 (n=23) |
| Protein Concentration (mol/µm$^2$) Mean ± SEM | 322 ± 9 (n=13) | 520 ± 13 (n=31) | 702 ± 9 (n=26) | 828 ± 6 (n=22) | 970 ± 12 (n=36) | 1283 ± 18 (n=37) | 1980 ± 70 (n=23) |
| Protein Brightness (KHz/mol) Mean ± SEM | 0.26 ± 0.07 (n=13) | 0.44 ± 0.13 (n=31) | 0.44 ± 0.09 (n=26) | 0.43 ± 0.6 (n=22) | 0.45 ± 0.05 (n=36) | 0.46 ± 0.05 (n=37) | 0.72 ± 0.17 (n=23) |
| Multimerization state Mean ± SEM | 11 ± 4 (n=13) | 18 ± 7 (n=31) | 22 ± 5 (n=26) | 19 ± 3 (n=22) | 20.0 ± 2.6 (n=36) | 20.4 ± 2.4 (n=37) | 33 ± 9 (n=23) |

**(v) Fit parameters connected to Figure S10A**

| **Parameter** | **DOPC** | **DOPC+DOPS** (9:1 molar ratio) | **DOPC+PI(4)P** (9:1 molar ratio) | **DOPC+PI(4,5)P2** (9:1 molar ratio) |
|---|---|---|---|---|
| *a* Mean ± SEM | 20 ± 21 (n=5) | 1.3 ± 1.5 (n=7) | 7 ± 3 (n=5) | 1.2 ± 1.1 (n=7) |
| *b* Mean ± SEM | 0.00 ± 0.16 (n=5) | 0.38 ± 0.18 (n=7) | 0.26 ± 0.06 (n=5) | 0.4 ± 0.13 (n=7) |

**Table S8:**

**(i) HCC parameters measured by lsFCS on GUVs related to Figure 4**

| **Parameter** | **DOPC GUV** | **DOPC+DOPS GUV** (9:1 molar ratio) | **DOPC+PI(4)P GUV** (9:1 molar ratio) | **DOPC+PI(4,5)P2 GUV** (9:1 molar ratio) |
|---|---|---|---|---|
| Protein Intensity (KHz) Mean ± SD | 0.6 ± 0.4 (n=4) | 3.8 ± 2 (n=51) | 2.8 ± 1.4 (n=38) | 2.9 ± 1.5 (n=38) |
| Protein Brightness (KHz/mol) Mean ± SD | | 0.10 ± 0.07 (n=50) | 0.10 ± 0.08 (n=30) | 0.10 ± 0.05 (n=35) |
| $D_{Protein}$ (μm$^2$/s) Mean ± SD | | 2.4 ± 1.8 (n=50) | 2.0 ± 1.8 (n=31) | 2.4 ± 1.6 (n=35) |

**(ii) Protein concentration vs. multimerization state on DOPC+DOPS (9:1) GUVs related to Figure S15**

| **Parameter** | **Point 1** | **Point 2** | **Point 3** | **Point 4** | **Point 5** |
|---|---|---|---|---|---|
| Protein Intensity (KHz) Mean ± SEM | 1.37 ± 0.17 (n=10) | 2.31 ± 0.06 (n=12) | 3.23 ± 0.12 (n=12) | 4.84 ± 0.19 (n=5) | 7.0 ± 0.4 (n=11) |
| Protein Concentration (mol/μm$^2$) Mean ± SEM | 23.60 ± 2.86 (n=10) | 39.800 ± 1.059 (n=12) | 55.70 ± 2.13 (n=12) | 83 ± 3 (n=5) | 120 ± 7 (n=11) |
| Protein Brightness (KHz/mol) Mean ± SEM | 0.068 ± 0.018 (n=10) | 0.09 ± 0.03 (n=12) | 0.094 ± 0.016 (n=12) | 0.12 ± 0.03 (n=5) | 0.136 ± 0.008 (n=11) |
| Multimerization state Mean ± SEM | 1.69 ± 1.50 (n=10) | 3.18 ± 2.65 (n=12) | 3.82 ± 1.33 (n=12) | 6.19 ± 2.73 (n=5) | 7.3500 ± 0.6813 (n=11) |

**(iii) Protein concentration vs. multimerization state on DOPC+PI(4)P (9:1) GUVs related to Figure S15**

| **Parameter** | **Point 1** | **Point 2** | **Point 3** | **Point 4** |
|---|---|---|---|---|
| Protein Intensity (KHz) Mean ± SEM | 1.24 ± 0.09 (n=6) | 2.06 ± 0.08 (n=9) | 2.90 ± 0.05 (n=5) | 9.0 ± 2.6 (n=10) |
| Protein Concentration (mol/μm$^2$) Mean ± SEM | 21.4 ± 2.58 (n=6) | 35.4 ± 1.32 (n=9) | 50 ± 1 (n=5) | 155 ± 40 (n=10) |
| Protein Brightness (KHz/mol) | 0.082 ± 0.019 (n=6) | 0.10 ± 0.03 (n=9) | 0.13 ± 0.04 (n=5) | 0.105 ± 0.021 (n=10) |

| | | | | |
|---|---|---|---|---|
| Mean ± SEM | | | | |
| Multimerization state<br>Mean ± SEM | 2.87 ± 1.55<br>(n=6) | 4.65 ± 2.81<br>(n=9) | 6.650 ± 3.078<br>(n=5) | 4.74 ± 1.74<br>(n=10) |

#### (iv) Protein concentration vs. multimerization state on DOPC+PI(4,5)P2 (9:1) GUVs related to Figure S15

| Parameter | Point 1 | Point 2 | Point 3 | Point 4 |
|---|---|---|---|---|
| Protein Intensity (KHz)<br>Mean ± SEM | 1.437 ± 0.020<br>(n=5) | 1.88 ± 0.08<br>(n=11) | 2.86 ± 0.12<br>(n=10) | 4.9 ± 0.4<br>(n=5) |
| Protein Concentration (mol/μm$^2$)<br>Mean ± SEM | 24.80 ± 0.34<br>(n=5) | 32.40 ± 1.43<br>(n=11) | 49.30 ± 2.08<br>(n=10) | 85 ± 7<br>(n=5) |
| Protein Brightness (KHz/mol)<br>Mean ± SEM | 0.071 ± 0.019<br>(n=5) | 0.087 ± 0.011<br>(n=11) | 0.094 ± 0.021<br>(n=10) | 0.128 ± 0.019<br>(n=5) |
| Multimerization state<br>Mean ± SEM | 1.918 ± 1.590<br>(n=5) | 3.22 ± 0.88<br>(n=11) | 3.84 ± 1.73<br>(n=10) | 6.660 ± 1.608<br>(n=5) |

#### v) Fit parameters for Figure S15

| **Parameter** | **DOPC+DOPS** (9:1 molar ratio) | **DOPC+PI(4)P** (9:1 molar ratio) | **DOPC+PI(4,5)P2** (9:1 molar ratio) |
|---|---|---|---|
| *a*<br>Mean ± SEM | 0.03650 ± 0.02019<br>(n=5) | 1.214 ± 1.320<br>(n=4) | 0.04690 ± 0.03104<br>(n=4) |
| *b*<br>Mean ± SEM | 1.077 ± 0.115<br>(n=5) | 0.239 ± 0.249<br>(n=4) | 1.078 ± 0.159<br>(n=4) |

**Table S9: HCC and Δ68 data connected to Figure S11**

| **Protein** | **Protein Brightness (KHz/mol) Mean ± SD** |
|---|---|
| HCC | 0.700 ± 0.500<br>(n=5) |
| Δ68 | 0.120 ± 0.0300<br>(n=15) |

**Table S10: PL *D* data in presence and absence of HCC related to Figures 5 and S16C**

| **Parameter** | **DOPC LD** (*pMMA*) | **DOPC+DOPS LD** (9:1 molar ratio) (*pMMA*) | **DOPC+PI(4)P LD** (9:1 molar ratio) (*pMMA*) | **DOPC+PI(4,5)P2 LD** (9:1 molar ratio) (*pMMA*) |
|---|---|---|---|---|
| $D_{PL}$ (µm$^2$/s) (-) HCC Mean ± SD | 7.7 ± 0.6 (n=29) | 6.6 ± 1.4 (n=48) | 5.6 ± 1.4 (n=45) | 5.3 ± 1.2 (n=45) |
| $D_{PL}$ (µm$^2$/s) (+) HCC Mean ± SD | 0.300 ± 0.205 (n=13) | 0.27 ± 0.18 (n=14) | 0.6 ± 0.4 (n=34) | 0.9 ± 0.9 (n=48) |
| $D_{PL/TF\text{-}PC}$ (µm$^2$/s) (-) HCC Mean ± SD | 3.2 ± 1.8 (n=48) | 3.1 ± 1.8 (n=29) | 3.8 ± 1.1 (n=38) | 4.5 ± 2.1 (n=37) |
| $D_{PL/TF\text{-}PC}$ (µm$^2$/s) (+) HCC Mean ± SD | 1.3 ± 0.8 (n=32) | 2.0 ± 1.2 (n=39) | 1.1 ± 1.2 (n=57) | 1.5 ± 1.3 (n=34) |
